# CXCL4 signaling and gene induction in human monocytes

**DOI:** 10.1101/2022.10.26.513860

**Authors:** Chao Yang, Ruoxi Yuan, Bikash Mishra, Richard D. Bell, Yaxia Zhang, Yong Du, Marie Dominique Ah Kioon, Franck J. Barrat, Lionel B. Ivashkiv

## Abstract

The chemokine CXCL4 activates myeloid cells and contributes to the pathogenesis of inflammatory and fibrotic diseases. One mechanism of CXCL4 action is binding of nucleic acids to promote their internalization and activation of endosomal TLRs. However, the signaling pathways and receptors that mediate myeloid cell responses to CXCL4 alone are not well characterized. Here, we report that in primary human monocytes, CXCL4 activated NF-κB and a TBK1-JNK signaling axis that drive the expression of inflammatory, fibrotic and neutrophil chemokine genes, and also RIPK3-dependent necroptosis. Surprisingly, six distinct lines of evidence targeting TLR4 expression and function suggested a role for TLR4 in CXCL4 responses. However, we were not able to completely dissect the contributions of CXCL4 alone to the observed results versus a contribution from endotoxin contamination. Our findings suggest that the interactions between CXCL4 and TLR4 merit further study.

## Main

Chemokine (C-X-C motif) ligand 4 (CXCL4), also known as platelet factor 4 (PF4), is a major product of platelets^1^, and large amounts of CXCL4 are released at sites of tissue injury when platelets are activated and degranulate to initiate the coagulation process. CXCL4 is also expressed by immune cells at sites of inflammatory pathology, such as plasmacytoid dendritic cells (pDCs) in fibrotic skin lesions in systemic sclerosis (SSc)^2, 3^ and macrophages in inflamed joint tissues in rheumatoid arthritis (RA)^4^. Accordingly, high levels of CXCL4 are found in skin wounds and are associated with multiple inflammatory conditions, including atherosclerosis, inflammatory bowel disease (IBD), SSc, and RA^2, 4, 5, 6^. CXCL4 can be present at concentrations as high as 10 μg/ml at sites of inflammation in IBD and 0.5 μg/ml in serum of SSc patients^2, 7, 8^. CXCL4 has been strongly implicated in inflammatory and related pro-fibrotic processes in SSc, primary myelofibrosis, and models thereof, where it can act on pDCs and endothelial, fibroblastic and myeloid cells^2, 3, 9, 10, 11^.

CXCL4 has wide-ranging effects on myeloid cells including induction of reactive oxygen intermediates and inflammatory cytokines and chemokines^1, 12^, macrophage differentiation^13^ and polarization of conventional DCs into a pro-fibrotic phenotype that is hyper-responsive to stimulation with ligands of nucleic acid-sensing endosomal Toll-like receptors (TLR3 and TLR8) ^3, 9, 14, 15^. The mechanism by which CXCL4 stimulates myeloid cells is not well understood; myeloid cells do not express CXCR3, the only known chemokine receptor for CXCL4, and myeloid cell activation by CXCL4 is independent of chemokine receptors and their associated G protein-mediated signaling pathways^1, 16^. One recently described mechanism by which CXCL4 activates cells independent of chemokine receptors is assembly of CXCL4 with “self” or microbial nucleic acids (NAs) into nanoparticles^1, 12, 17, 18^, which facilitates internalization of bound NAs to augment activation of endosomal NA-sensing Toll-like receptors (TLRs) such as TLR8/9 in pDCs and macrophages^7, 12, 18^. In human macrophages, formation of CXCL4-NA nanoparticles contributes to synergistic activation of inflammatory genes and of the NLRP3 inflammasome when cells are stimulated simultaneously with CXCL4 and a single stranded RNA TLR8 ligand^12^.

Several groups including ours have reported activation of inflammatory signaling pathways and chromatin remodeling at inflammatory genes in myeloid cells after stimulation with CXCL4 alone, in the absence of exogenous nucleic acids^1, 10, 12, 14, 18^. This suggests that CXCL4 can also activate myeloid cells independently of its NA-chaperone properties. However, mechanistic understanding of myeloid cell activation by CXCL4 has been limited over several decades of investigation by the lack of a high affinity cognate receptor that mediates CXCL4 signaling events. Instead, CXCL4 binds to cells via low affinity charge-charge interactions with cell surface proteoglycans (PGs)^17^. In accord with a well-established model whereby PGs serve as co-receptors that facilitate interactions of various growth factors, chemokines, and cytokines with their receptors^19^, PG-mediated presentation of CXCL4 to cell surface receptors such as LDLR and αMβ2 integrins has been proposed to mediate cell activation^20, 21^. Binding of CXCL4 to αMβ2 (also called Mac-1) changes cell morphology, shifts αMβ2 to an intracellular location co-localized with actin, and increases cell migration and phagocytosis^21^. However, the receptors and signaling pathways that mediate inflammatory activation of macrophages by CXCL4 are not well characterized.

Cell responses to bacterial lipopolysaccharide (LPS) are mediated by TLR4. TLR4 is expressed on the plasma membrane as part of a heterodimer with Myeloid Differentiation Protein-2 (MD-2), which is required for LPS sensing and cell activation. TLR4 forms only weak and minimal contacts with LPS, and effective activation of TLR4 by LPS, which occurs at picomolar concentrations, requires MD-2 and additional LPS-binding proteins such as CD14 that act upstream of and sensitize responses to TLR4 (reviewed in ^22^). CD14 binds and transfers LPS to MD-2, which results in TLR4 dimerization and activation of Myd88-mediated NF-κB and MAPK signaling. TLR4 subsequently traffics to endosomes, where its signaling is mediated by a distinct adaptor TRIF that, in addition to NF-κB and MAPKs, activates transcription factor IRF3 and downstream IFN responses. A role for TLR4 has also been proposed in the activation of macrophages by various endogenous proteins and proteoglycans that are released during cell death or degradation of extracellular matrix (termed DAMPs or alarmins) (reviewed in ^23, 24^).

Interactions between some DAMPs and TLR4 have been detected in the nanomolar range and by co-immunoprecipitation experiments, and one model proposes that these DAMPs work as ligands and direct agonists of TLR4. A different model proposes that these DAMPs may instead act by binding LPS and presenting it to TLR4/MD-2, thus sensitizing macrophages to respond to low LPS concentrations, similar to the function of CD14^23^. In this paradigm, LPS-binding DAMPs function as TLR4-sensitizing molecules, and this sensitization function can also occur in vivo, where DAMPs can bind LPS derived from the microbiome, or from bacteria present during bacteremia or at wound sites.

In our previous study of cooperation between CXCL4 and TLR8 in macrophage activation, we found that CXCL4 alone substantially activated NF-κB and MAPK signaling and weakly activated TANK-binding kinase 1 (TBK1), whereas TLR8 strongly activated interferon-regulated factors (IRFs) and synergized with CXCL4 to potently activate TBK1^12^. In the current study, we investigated mechanisms by which CXCL4 alone activates human monocytes. We found that NF-LJB and a TBK1-JNK signaling axis are important for induction of inflammatory and tissue repair/fibrotic genes by CXCL4, and CXCL4 also induced cell death in a RIPK3- and MLKL-dependent manner. Surprisingly, induction of inflammatory and fibrotic genes and related signaling pathways was strongly dependent on TLR4, but CXCL4 only minimally induced expression of interferon-stimulated genes (ISGs) and IL-12 family genes compared to the canonical TLR4 agonist LPS. We were not able to completely dissect the contributions of CXCL4 alone to the observed results versus a contribution from endotoxin contamination and whether TLR4 becomes activated or signals in a noncanonical manner in the presence of CXCL4. These results identify mechanisms by which CXCL4 induces and regulates macrophage activation.

## Results

### CXCL4 activates inflammatory and tissue repair pathways in human monocytes

We used a transcriptomic approach to gain insight into how CXCL4 activates human monocytes. Our previous work^12^ showed that CXCL4 rapidly but transiently induces expression of inflammatory genes such as *TNF* and *IL6*, with a return to near baseline 6 hours after CXCL4 stimulation. Here, we analyzed the transcriptomics of the CXCL4 response at 1 h and 3 h in human monocytes using RNA-seq (Extended Data Fig.1a). CXCL4 stimulation significantly (FDR < 0.05, fold change ≥ 2) upregulated expression of 1,771 genes and downregulated expression of 1,840 genes at 3 h (Fig. 1a). Various inflammatory and tissue repair/fibrotic genes were significantly upregulated, including *IL6*, *IL1B*, *TNF*, *CXCL1*, *PDGFA* and *PDGFB* (Fig. 1a). Significantly differentially expressed genes (DEGs) (4,004; FDR < 0.05, fold change ≥ 2) clustered into 4 groups based on pattern of expression (Fig. 1b). Genes in groups III and IV were early response-genes whose expression changed at 1 h after CXCL4 stimulation while changes in expression of genes in groups I and II were observed at 3 h (Fig. 1b). Gene-ontology (GO) and Ingenuity Pathway Analysis (IPA) analysis^25^ of genes regulated by CXCL4 at 3 h showed most significant enrichment of cytokine-mediated signaling, immune response, apoptotic and wound healing/fibrosis pathways (Fig. 1c and 1d). Gene Set Enrichment Analysis (GSEA)^26, 27^ further supported activation of wound healing and related TGF-β1 signaling and revealed enrichment of genes in myeloid cell differentiation and JNK signaling pathways (Fig. 1e and Extended Data Fig. 1c). CXCL4-mediated induction of representative inflammatory, fibrotic, and wound healing genes is shown in heatmaps (Fig. 1f and Extended Data Fig. 1d - f). Notably, CXCL4 induced expression of neutrophil chemokine genes CXCL1, CXCL2 and CXCL3 that are relevant for neutrophil infiltration into wound sites. CXCL4 still significantly induced *TNF* expression when any endogenous extracellular DNA and RNA were depleted from cultures (Extended Data Fig. 2a), supporting that CXCL4 can activate cells independently of forming complexes with NAs. Overall, the data show that CXCL4 broadly activates inflammatory and profibrotic gene expression, and induces neutrophil chemokine genes, in human monocytes.

**Figure 1.**
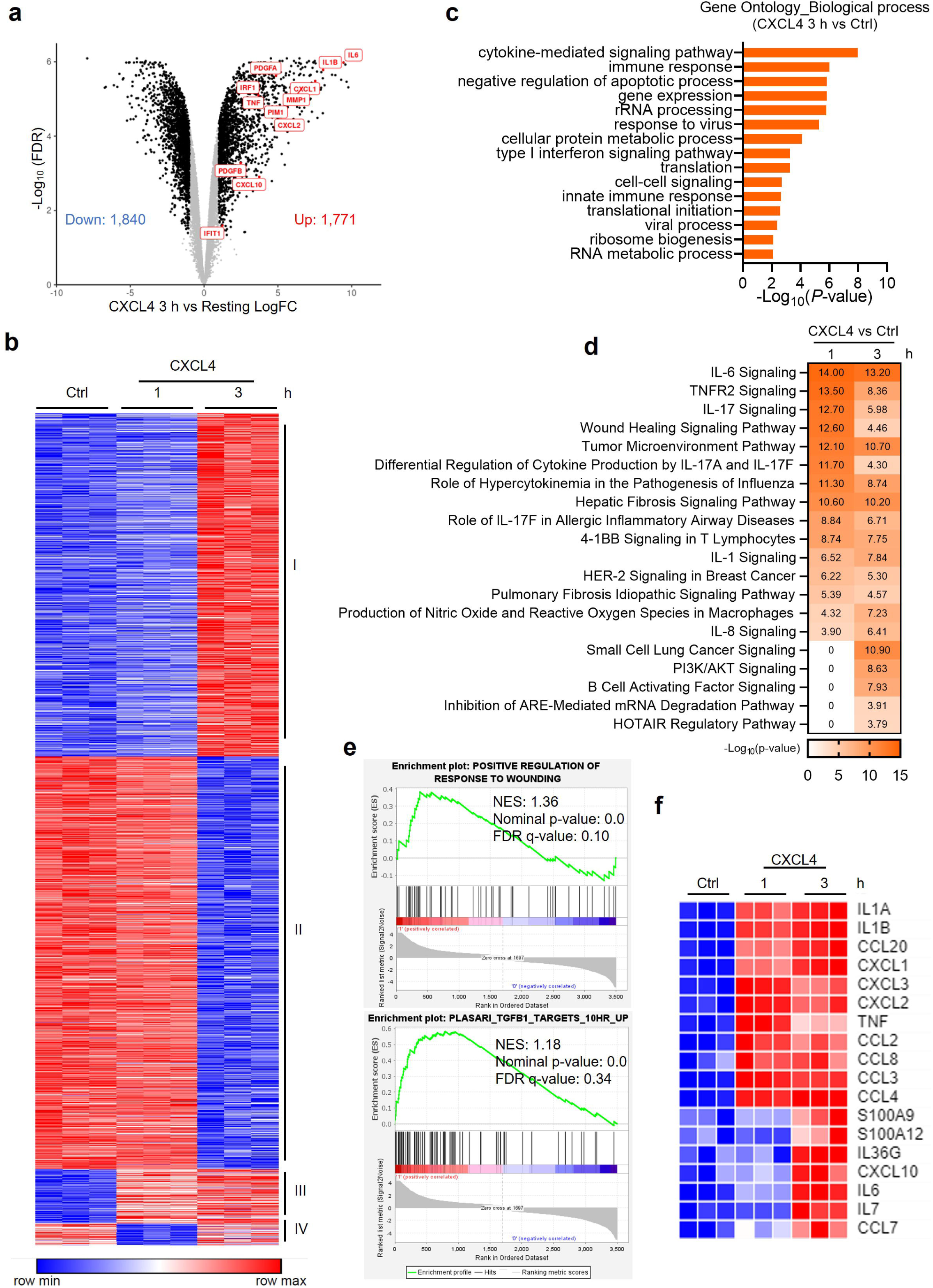
(a) Volcano plot of DEGs regulated by CXCL4 (10 µg/ml) after 3 h treatment of human primary macrophages. (b) k-means clustering of the DEGs regulated in human primary macrophages by CXCL4 after 1 h and 3 h treatments (n = 3 independent donors). (c) Gene Ontology analysis using program InnateDB of DEGs regulated in human primary macrophages by CXCL4 after 3 h treatment. (d) IPA analysis of DEGs regulated in human primary macrophages by CXCL4 after 1 h and 3 h treatments. (e) GSEA analysis of DEGs after CXCL4 3 h treatment. (f) Heatmap of inflammatory genes regulated by CXCL4.

### NF-**κ**B and a TBK1-JNK axis drive inflammatory, fibrotic and neutrophil chemokine gene expression

CXCL4 alone substantially activates NF-κB and MAPKs p38 and ERK, and also activates TBK1^12^. We next investigated the role of these signaling molecules in induction of representative inflammatory and fibrotic genes, and the neutrophil chemokines CXCL1 and CXCL2. Activation of NF-κB was dependent on IKKα/β and TAK1 (Extended Data Fig. 3a), and as expected, inhibition of IKKα/β suppressed CXCL4-induced expression of canonical inflammatory NF-κB target genes *IL6*, *TNF*, *IL1B* and *IRF1* (Extended Data Fig. 3b). Strikingly, induction of representative wound healing/fibrotic genes *PDGFA* and *PDGFB* was essentially completely blocked by NF-κB inhibition, while induction of *CXCL1* and *CXCL2* was relatively resistant (Extended Data Fig. 3c). These results implicate NF-κB in the regulation of fibrotic in addition to inflammatory genes in CXCL4-stimulated monocytes.

We next confirmed activation of TBK1 by CXCL4 alone in an additional 3 donors (Fig. 2a). In human monocytes activation of TBK1 is coupled with induction of inflammatory NF-κB target genes^12, 28^. Accordingly, three different TBK1 inhibitors suppressed CXCL4-mediated induction of inflammatory genes *TNF* and *IL1B* (Fig. 2b). Induction of fibrotic genes *PDGFA*, *PDGFB* and *CXCL1* was suppressed, while *IL6* and *IRF1* were not affected (Fig. 2b and Extended Data Fig. 4a). As inhibition of TBK1 does not diminish NF-κB activation^12^, we tested whether TBK1 regulates MAPK signaling. In addition to activating ERK and p38^12^, CXCL4 activated MAPK JNK p46 and p54 isoforms, and this activation was suppressed when TBK1 was inhibited (Fig. 2c). CXCL4 also activated the upstream MAPK kinases MKK4/7 that phosphorylate and activate JNK in canonical MAPK signaling pathways^29^. However, CXCL4-induced MKK4/7 phosphorylation was not sensitive to TBK1 inhibitors (Extended Data Fig. 4b), suggesting TBK1 can activate JNK by a noncanonical pathway. In accord with this notion, JNK co-immunoprecipitated with TBK1 in unstimulated human monocytes, which suggests that TBK1 interacts with JNK even in the steady state; CXCL4 stimulation did not consistently increase the interaction of JNK with TBK1 (Fig. 2d and Extended Data Fig. 4c).

**Figure 2.**
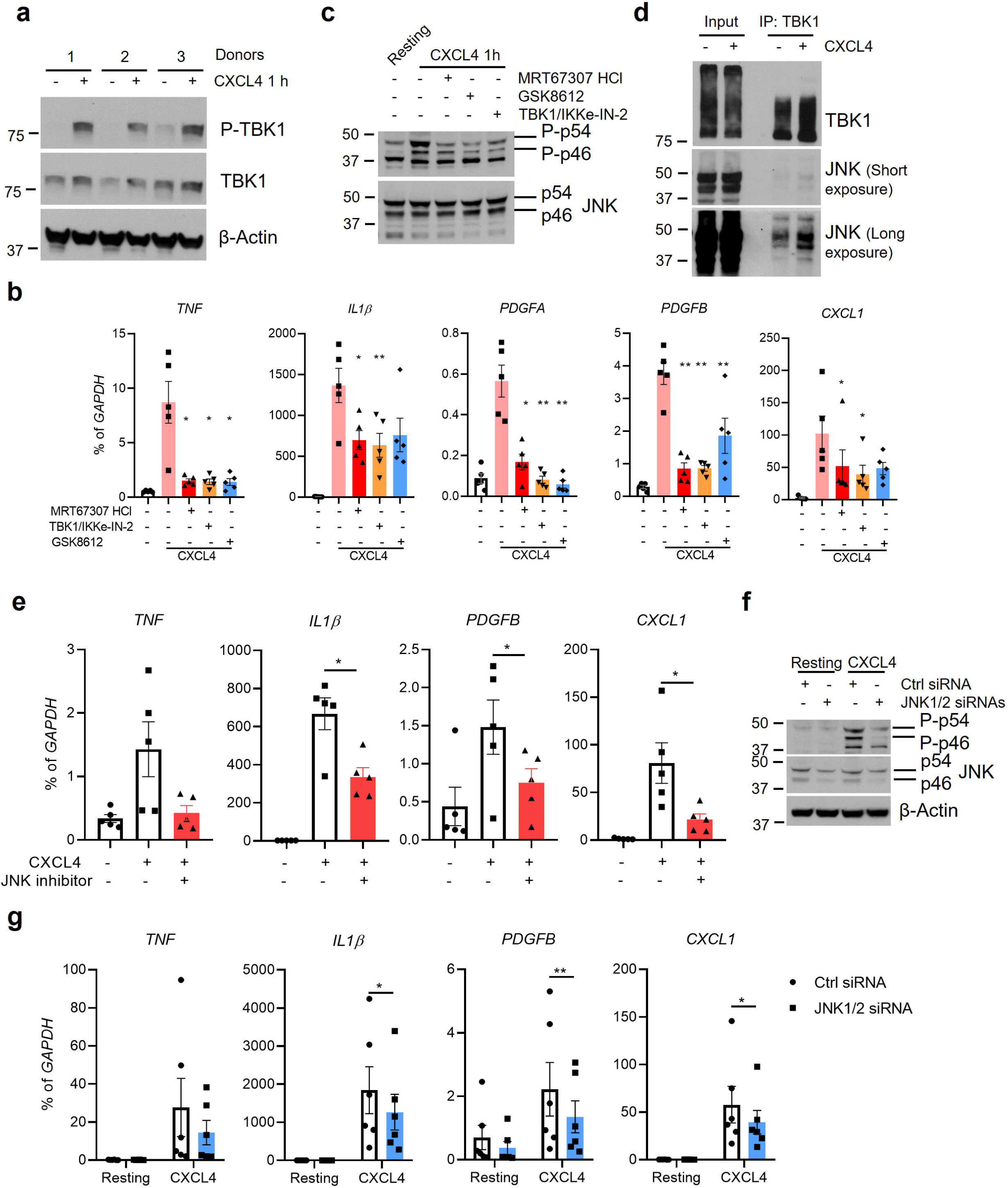
(a) Immunoblot of phospho-TBK1 and TBK1 with whole cell lysates of cells stimulated with CXCL4 for 1 h (n = 3 independent donors). (b) mRNA of *TNF*, *IL1*β, *PDGFA*, *PDGFB* and *CXCL1* measured by qPCR and normalized relative to *GAPDH* mRNA in cells pre-treated with TBK1/IKKε inhibitors MRT67307 HCl (10 µM), TBK1/IKKε-IN-2 (1 µM) or GSK8612 (50 µM) for 30 min and then stimulated with CXCL4 for 3 h (n = 5 independent donors). (c) Immunoblot of phospho-JNK (p54 and p46) and total JNK with whole cell lysates of cells pre-treated with TBK1/IKKε inhibitors MRT67307 HCl (10 µM), TBK1/IKKε-IN-2 (1 µM) or GSK8612 (50 µM) for 30 min and then stimulated with CXCL4 for 1 h (representative of 4 independent donors). (d) Immunoblot of TBK1 and JNK from whole cell lysates immunoprecipitated with TBK1 antibodies (representative of 5 independent donors). (e) mRNA of indicated genes measured by qPCR and normalized relative to *GAPDH* mRNA in cells pre-treated with JNK inhibitor SP600125 (10 µM) for 30 min and then stimulated with CXCL4 for 3 h (n = 5 independent donors). (f) Immunoblot of phospho-JNK and JNK using whole cell lysates 3 days after nucleofection of monocytes with control or JNK-specific siRNAs (representative of 3 independent donors). (g) mRNA of indicated genes was measured by qPCR and normalized relative to *GAPDH* mRNA after knockdown of JNK by siRNA for 3 days (n = 6 independent donors). Data are depicted as mean ± SEM (b, e, g). **p ≤ 0.01; *p ≤ 0.05 by One-way ANOVA (b, e) or Two-way ANOVA (g). Source data are provided as a Source Data file.

Next, we evaluated the contribution of JNK to CXCL4-induced gene expression. Inhibition of JNK significantly suppressed CXCL4-mediated induction of *TNF*, *IL1B, PDGFB* and *CXCL1* (Fig. 2e and Extended Data Fig. 4d). A role for JNKs in CXCL4-induced gene expression was corroborated using siRNA-mediated knockdown of both JNK1 and JNK2 (Fig. 2f, g and Extended Data Fig. 4e). The effects of inhibiting ERKs and p38 were more limited than inhibition of JNKs (Extended Data Fig. 5). Collectively, these data highlight the importance of a CXCL4-activated TBK1-JNK axis in induction of key inflammatory, fibrotic and neutrophil chemokine genes.

### CXCL4 activates STAT3 via IL-10 autocrine signaling

CXCL4 could activate STAT3, as assessed by phosphorylation at tyrosine 705 (Extended Data Fig. 6a). Inflammatory factors typically activate Jak-STAT signaling indirectly via induction of cytokines that function in an autocrine manner. In accord with this model, inhibition of cytokine secretion using Brefeldin A (BFA) abolished CXCL4-inducedSTAT3 activation. The STAT3-activating cytokine IL-10 was induced by CXCL4 (Extended Data Fig. 6b, c) and blockade of IL-10 function using IL-10Rα neutralizing antibody completely abolished CXCL4-induced STAT3 phosphorylation (Extended Data Fig. 6d). Although autocrine IL-10 downregulates inflammatory gene ^30, 31^, knockdown of STAT3 using siRNA (Extended Data Fig. 6e) had minimal effect on CXCL4-induced *IL6*, *TNF*, *IL1B*, and *IRF1* expression (Extended Data Fig. 6g), possibly because of low amounts of endogenous IL-10 or incomplete STAT3 knockdown. However, STAT3 knockdown significantly decreased induction of the known STAT target gene *PIM1* and of *PDGFA* (Extended Data Fig. 6f); in contrast, induction of *CXCL1* and *CXCL2* were increased. These results implicate STAT3 in CXCL4-mediated gene regulation and suggest that STAT3 can play a positive role in induction of fibrotic genes in addition to its established role in monocytes of suppressing inflammatory responses.

### CXCL4 activates a TRIF-utilizing receptor to induce RIPK3-dependent necroptosis

Only a relatively small subset of inflammatory factor receptors can activate RIPK3 and downstream necroptotic cell death - TNF receptor family members that contain a death domain and activate RIPK3 via RIPK1, and receptors that utilize the TRIF signaling adaptor (namely, TLR3 and TLR4) and can activate RIPK3 directly. To try to narrow down the receptor(s) that CXCL4 engages to induce inflammatory responses, we tested whether CXCL4 induces RIPK3-mediated necroptosis, which is mediated by downstream effector MLKL and requires inhibition of Caspase-8 to prevent inactivating cleavage of RIPK3^32^. Stimulation of monocytes with CXCL4 in the presence of the Caspase inhibitor Z-VAD-FMK (zVAD) resulted in a significant induction of cell death (Fig. 3a). CXCL4-induced cell death was significantly blocked by inhibition of MLKL using necrosulfonamide (NSA) (Fig. 3a) and by siRNA-mediated knockdown of RIPK3 or MLKL (Fig. 3b and Extended Data Fig. 7a). In contrast, knockdown of RIPK1 using siRNA did not reduce cell death (Fig. 3c and Extended Data Fig. 7b); the dispensability of RIPK1 makes upstream involvement of TNF family receptors less likely.

**Figure 3.**
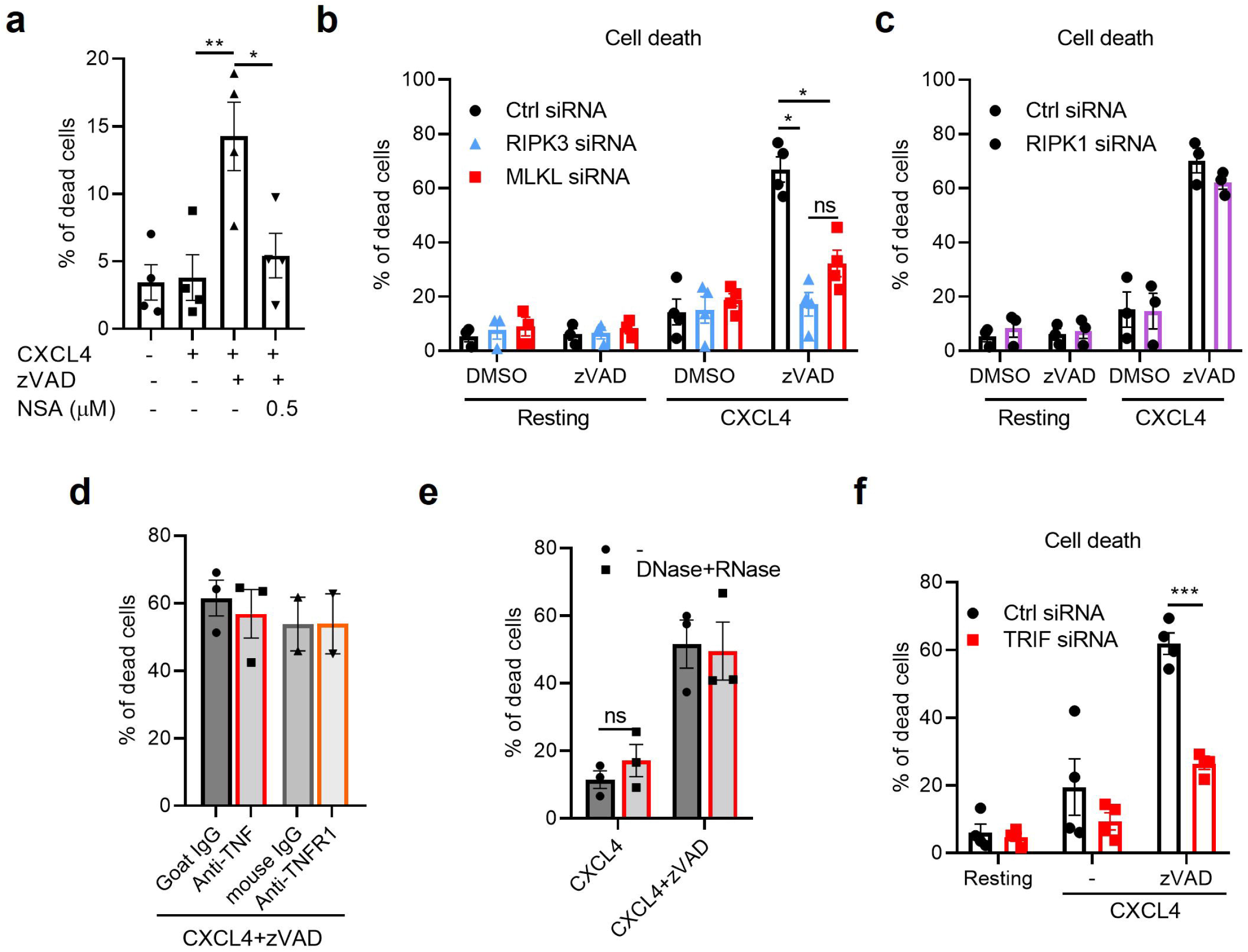
(a) Cell death assay of human macrophages stimulated with CXCL4 in the presence/absence of Caspase-8 inhibitor zVAD (50 µM) and necrosulfonamide (NSA, 0.5 µM) for 6 h (n = 4 independent donors). (b) Cell death assay performed 3 days after nucleofection of monocytes with control (Ctrl) siRNA, RIPK3 or MLKL specific siRNAs, and then stimulated with CXCL4 in the presence/absence of Caspase-8 inhibitor zVAD for 6 h (n = 4 independent donors). (c, f) Cell death assay performed 3 days after nucleofection of monocytes with control (Ctrl) siRNA, RIPK1 (c) or TRIF (f) specific siRNAs, and then stimulated with CXCL4 in the presence/absence of Caspase-8 inhibitor zVAD for 6 h (n = 3 (c) or 4 (f) independent donors). (d, e) Cell death assay in cells pretreated with TNF (n = 4 independent donors) or TNFR1 (n = 2 independent donors) neutralizing antibodies (d), or with DNase I (1000 U) + RNase A (50 ug/ml) (n = 3 independent donors) (e) for 1 h and then stimulated with CXCL4 in the presence of zVAD for 6 h. Data are depicted as mean ± SEM. ***p ≤ 0.001; **p ≤ 0.01; *p ≤ 0.05 by One-way ANOVA (a) or Two-way ANOVA (b, e, f). Source data are provided as a Source Data file.

Furthermore, neutralizing TNF antibodies and TNFR1 blocking antibodies did not reduce CXCL4-induced cell death (Fig. 3d), arguing against a role for autocrine TNF. Additionally, degradation of extracellular nucleic acids in culture supernatants did not affect CXCL4-induced cell death (Fig. 3e), suggesting that CXCL4 did not induce cell death by promoting NA internalization. Finally, we tested the role of the signaling adaptor TRIF, which activates RIPK3 and contributes to TLR3/4-mediated necroptosis^33, 34, 35^. siRNA-mediated knock down of TRIF almost completely abolished CXCL4-induced necroptosis (Fig. 3f). These data suggest that CXCL4 activates a TRIF-utilizing receptor.

### CXCL4 minimally activates IFN responses and IL-12 genes and is resistant to IFN priming in a gene-specific manner

To our knowledge, the only receptors that utilize TRIF for signaling are TLR3 and TLR4^22^, and TRIF links these receptors to strong IFN responses mediated by induction of autocrine IFN-β that binds to its receptor IFNAR to activate Jak-STAT signaling and expression of interferon-stimulated genes (ISGs). We thus tested whether CXCL4 similarly induces an IFN response and as a comparator used the canonical TLR4 agonist LPS, which activates a robust IFN autocrine loop via TRIF-mediated signaling. CXCL4 induced expression of inflammatory genes *TNF* and *IL1B* comparably to LPS (5 ng/ml) (Fig. 4a). In striking contrast, CXCL4 minimally induced expression of ISGs *CXCL10* and *IFIT1* (Fig. 4a); the amounts of CXCL10 and IFIT1 mRNA induced by CXCL4 were at least two orders of magnitude lower than those induced by LPS. Induction of *IFNB* mRNA by CXCL4 was also lower than induction by LPS (Fig. 4a and 6b).

**Figure 4.**
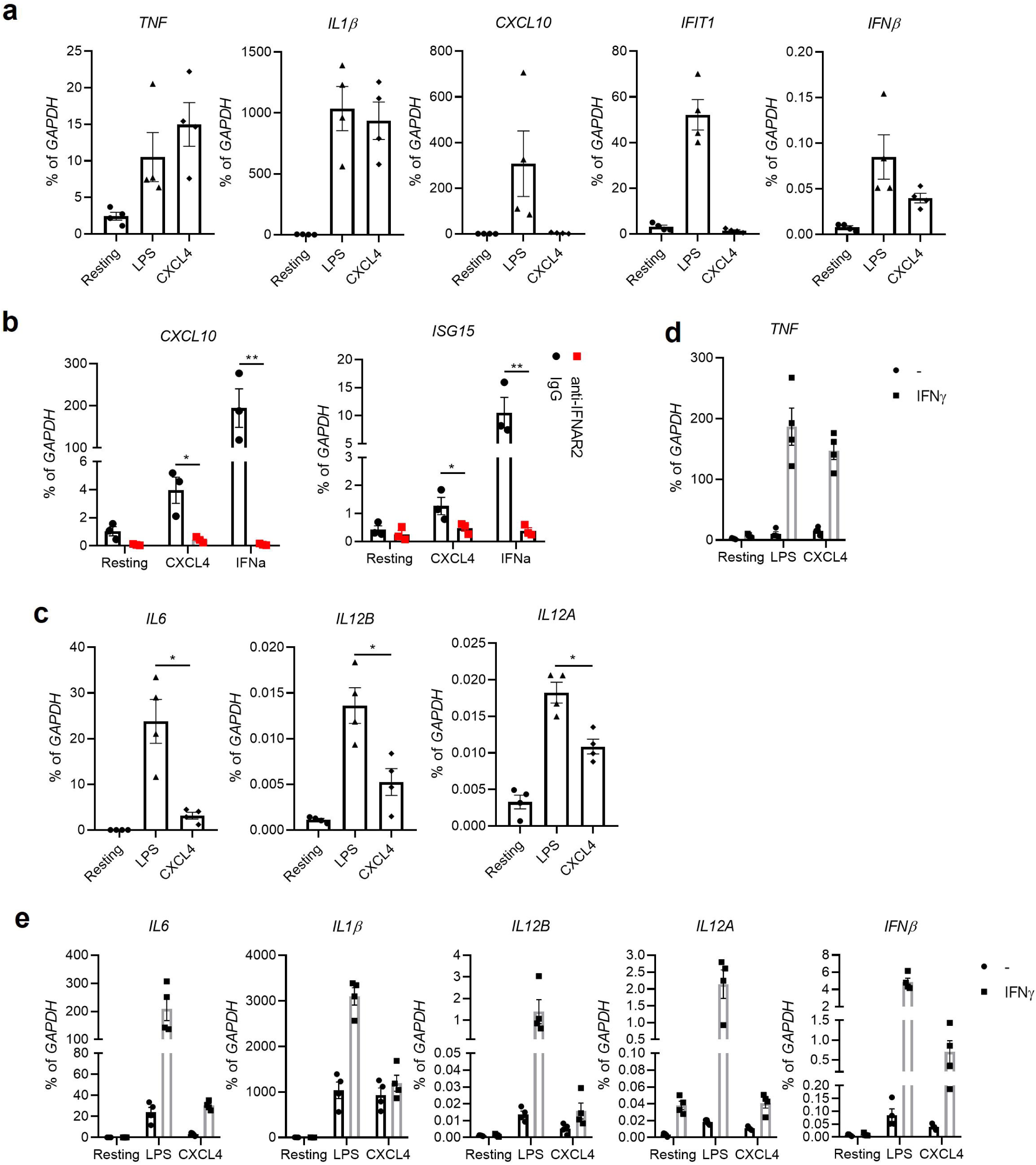
(a, c) mRNA of indicated genes measured by qPCR and normalized relative to *GAPDH* mRNA in cells stimulated with LPS (5 ng/ml) or CXCL4 for 3 h (n = 4 independent donors).(b) mRNA of *CXCL10* and *ISG15* measured by qPCR and normalized relative to *GAPDH* mRNA in cells pre-treated with IFNAR2 neutralizing antibody (10 µg/ml) for 1 h and then stimulated with CXCL4 for 3 h (n = 3 independent donors). (d, e) mRNA of indicated genes measured by qPCR and normalized relative to *GAPDH* mRNA in cells pretreated with IFN_γ_ (100 U/ml) for 24 h and then stimulated with LPS (5 ng/ml) or CXCL4 for 3 h (n = 4 independent donors). The data in a, c, d, and e are derived from the same donors and the resting, LPS and CXCL4 conditions in a, c, d and e represent the same samples; data are depicted separately to enhance visualization of the comparison of samples performed in the absence of IFN-γ in a and c. Data are depicted as mean ± SEM. ***p ≤ 0.001; **p ≤ 0.01; *p ≤ 0.05 by Two-way ANOVA. Source data are provided as a Source Data file.

Induction of ISGs by CXCL4 was mediated by autocrine IFNs as ISG induction was abolished by blocking antibodies to IFNAR2 (Fig. 4b), but induction of ISGs by CXCL4 was massively lower than induction of ISGs by exogenous IFN-α (Fig. 4b). Collectively these results indicate that although CXCL4 activates inflammatory gene expression comparably to LPS, CXCL4 only minimally induces an IFN response.

Another distinction between CXCL4 and TLR4 responses was weak induction of *IL6* and *IL12B* and *IL12A* (Fig. 4c). A core feature of the LPS response in human monocytes is that induction of NF-κB target genes like *IL6* and *IL12B* is increased by pretreatment of cells with IFN-γ (termed priming)^36^. As expected, IFN-γ priming strongly increased the amounts of *TNF, IL6*, *IL1B*, *IFNB*, *IL12B* and *IL12A* mRNA induced by LPS (Fig. 4d and e). IFN-γ priming also increased the amounts of *TNF* mRNA induced by CXCL4 (Fig. 4d). In striking contrast, the amounts of *IL6*, *IL1B* and *IFNB* mRNA induced by CXCL4 were minimally increased by IFN-γ priming, and induction of *IL12B* and *IL12A* genes remained refractory to CXCL4 stimulation even in IFN-γ-primed macrophages (Fig. 4e). Thus, the difference between CXCL4 and LPS in responsiveness to IFN-γ priming was gene-specific. However, as the effects of IFN-γ priming are context-dependent and can vary with state of differentiation of cells and concentrations of LPS used. As we observed preferential IFN-γ-mediated priming of *TNF* when low amounts of LPS were used (data not shown), these results do not definitively differentiate CXCL4 from LPS stimulation.

### CXCL4 signaling and gene activation diminish when TLR4 is blocked

Since CXCL4 activates a TRIF-dependent necroptosis pathway, we considered known receptors that use TRIF as an adaptor, TLR3 and TLR4, as candidates for mediating the effects of CXCL4. TLR3 is minimally expressed in human macrophages^37^ and stimulation of human macrophages with the TLR3 ligand poly (I:C) does not induce gene expression^12^. Thus, despite the differences between CXCL4 and LPS responses described above, we tested whether TLR4 is involved in CXCL4 responses in human macrophages. The small molecule inhibitor of TLR4 signaling CLI-095 that binds to the TLR4 intracellular domain abolished both CXCL4- and LPS-induced expression of *TNF*, *IL1*β and *IL6* (Extended Data Fig. 8a). As a specificity control, CLI-095 did not affect gene induction by TLR2 ligand Pam3CSK4 (Extended Data Fig. 8b). In our recent study, we showed that the very low amount of LPS present in recombinant CXCL4 (1-2 pg/ml final concentration in experiments) does not have the ability to activate expression of inflammatory genes in human macrophages^12^, and we found similarly low concentrations of LPS in additional lots of CXCL4 (Extended Data Fig. 9); however, we also found variable concentrations of contaminating LPS in various lots of CXCL4 purchased from different vendors (Extended Data Fig. 10a). CXCL4 completely lost its activity upon heat denaturation of proteins (Fig. 5a and Extended Data Fig. 8c). Although LPS is typically considered to be heat-resistant, we found that the activity of low amounts of LPS was diminished by heating, and thus this result on its own is insufficient to exclude that low amounts of contaminating endotoxin contributed to the observed results (Extended Data Fig. 10b).

**Figure 5.**
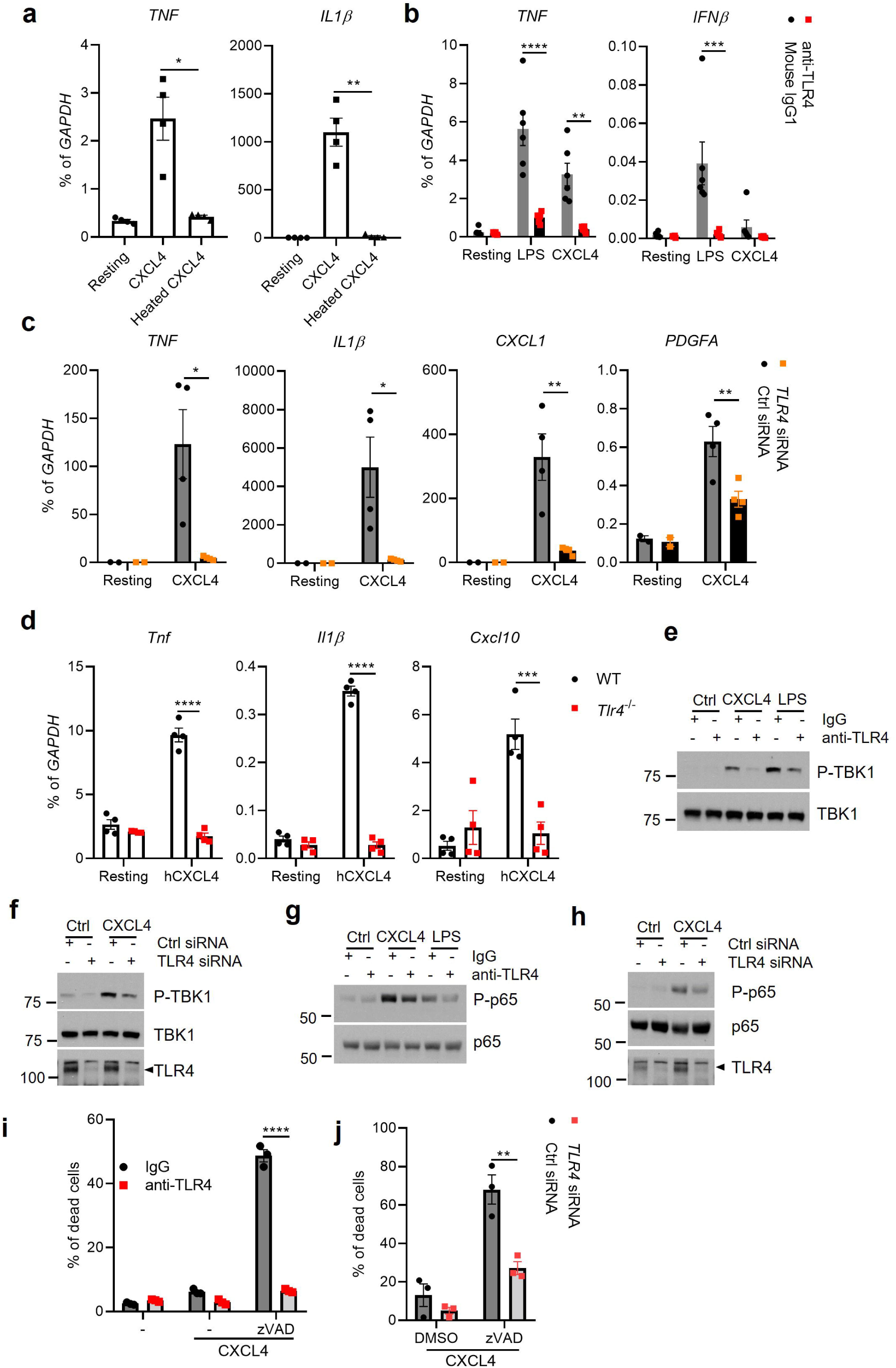
(a) mRNA of *TNF* and *IL1B* measured by qPCR and normalized relative to *GAPDH* mRNA in cells treated with CXCL4 or heat denatured CXCL4 for 3 h (n = 4 independent donors). (b) mRNA of *TNF* and *IFNB* measured by qPCR and normalized relative to *GAPDH* mRNA in cells pre-treated with TLR4 neutralizing antibody (5 µg/ml) for 1 h and then stimulated with CXCL4 or LPS (5 ng/ml) for 3 h (n = 6 independent donors). (c) mRNA of indicated genes was measured by qPCR and normalized relative to *GAPDH* mRNA in cells after knockdown of TLR4 by specific siRNA for 3 days and then stimulation with CXCL4 for 3 h (n = 4 independent donors). (d) mRNA of indicated genes was measured by qPCR and normalized relative to *GAPDH* mRNA in WT and *Tlr4*^-/-^ BMDMs stimulated with CXCL4 for 3 h (n = 4 independent experiments). (e, f) Immunoblot of phospho-TBK1 and TBK1 with whole cell lysates of cells pre-treated with TLR4 neutralizing antibody (e) for 1 h or after knockdown of TLR4 by specific siRNA (f) for 3 days and then stimulated with CXCL4 for 3 h (representative of 4 independent donors for both e and f). (g, h) Immunoblot of phospho-p65 and p65 with whole cell lysates of cells pre-treated with TLR4 neutralizing antibody (g) for 1 h or after knockdown of TLR4 by specific siRNA (h) for 3 days and then stimulated with CXCL4 for 3 h (representative of 4 independent donors for both g and h). (i, j) Cell viability in cells pre-treated with TLR4 neutralizing antibody (i) for 1 h or after knockdown of TLR4 by specific siRNA (j) for 3 days and then stimulation with CXCL4 in the presence/absence of zVAD for 3 h (n = 3 independent donors for both i and j). Data are depicted as mean ± SEM (a - d, i and j). ****p ≤ 0.0001; ***p ≤ 0.001; **p ≤ 0.01; *p ≤ 0.05 by One-way ANOVA (a) or Two-way ANOVA (b - d, i and j). Source data are provided as a Source Data file.

**Figure 6.**
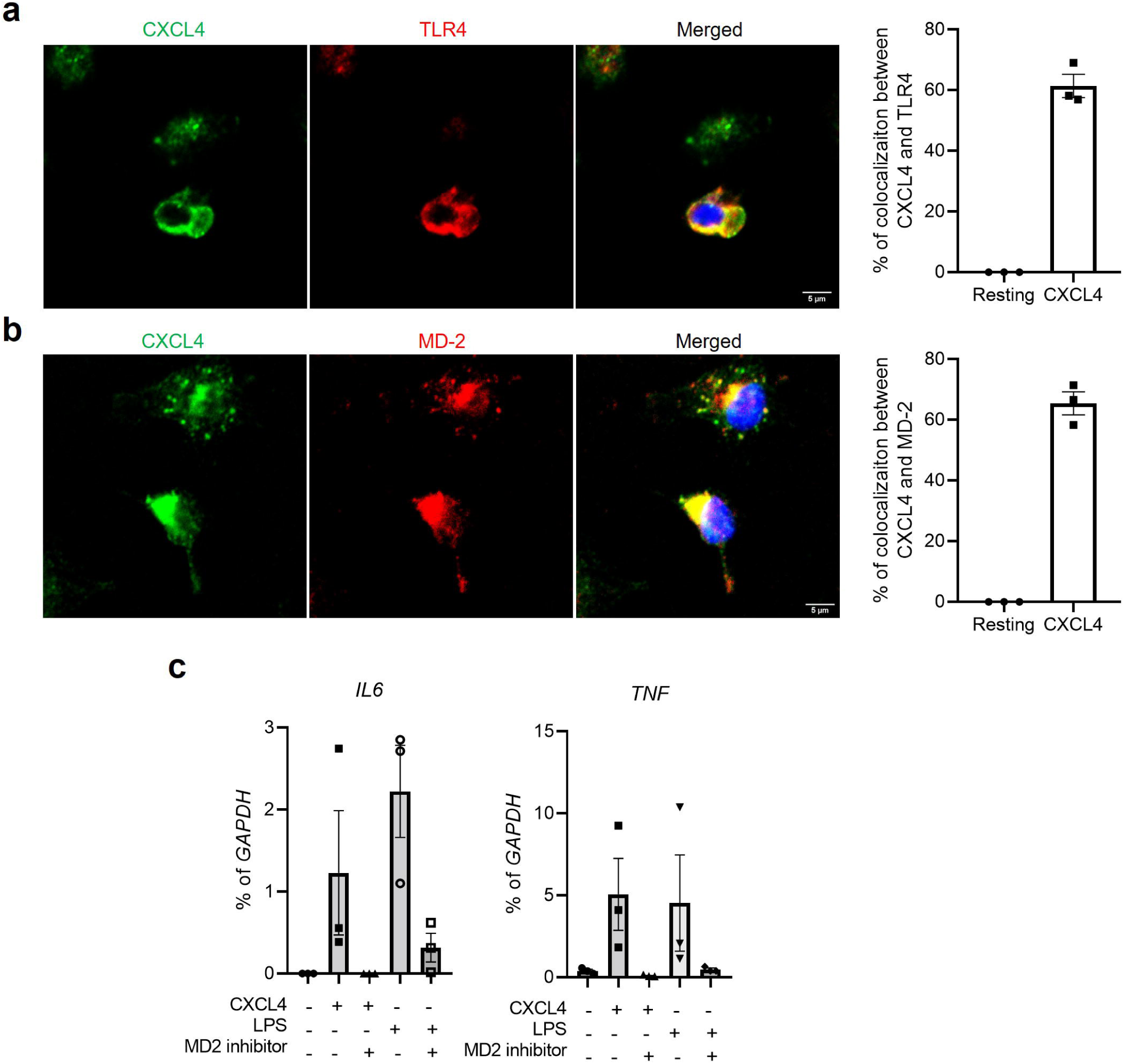
(a, b) Confocal microscopy images of CXCL4 colocalization with TLR4 (a) and MD-2 (b) in human macrophages 30 min after treatment (representative images from 3 independent experiments with different donors for each TLR4 and MD-2). (c) mRNA of *IL6* and *TNF* measured by qPCR and normalized relative to *GAPDH* mRNA in cells pre-treated with MD-2 inhibitor MD2-IN-1 (10 µM) for 30 min and then stimulated with CXCL4 or LPS (5 ng/ml) for 3 h (n = 3 independent donors). Data are depicted as mean ± SEM (c). Source data are provided as a Source Data file.

To further investigate whether TLR4 plays a role in CXCL4 responses, we blocked TLR4 using a specific neutralizing antibody and found that TLR4 blockade nearly completely abolished induction of inflammatory, fibrotic and neutrophil chemokine genes by both CXCL4 and LPS (Fig. 5b and Extended Data Fig. 8d). Although induction of *IFNB* by CXCL4 was lower than induction by LPS, the effects of both stimuli were strongly suppressed by TLR4 blockade (Fig. 5b). Further, genetic knockdown of TLR4 using its specific siRNA corroborated the role of TLR4 in CXCL4-induced gene expression in human macrophages (Fig. 5c and Extended Data Fig. 8e). Finally, we stimulated *Tlr4*^-/-^ bone marrow-derived mouse macrophages (BMDMs) with human CXCL4 (hCXCL4) and found that deletion of *Tlr4* abolished CXCL4-induced gene expression (Fig. 5d). In accord with decreased gene induction, blockade of TLR4 using neutralizing antibodies or knockdown of TLR4 using siRNA suppressed CXCL4-induced TBK1 and NF-LJB activation (Fig. 5e-h), and also abolished CXCL4-induced necroptosis (Fig. 6i and j). These studies provide four independent lines of evidence implicating TLR4 in mediating macrophage responses to CXCL4.

### CXCL4 colocalizes with TLR4 and MD-2

Next, we investigated whether CXCL4 colocalized with TLR4. To do so, we treated human primary macrophages with CXCL4 for 30 min and the cells were fixed and stained with anti- CXCL4 and anti-TLR4 antibodies for immunofluorescence analysis. Addition of CXCL4 altered TLR4 staining to a more discrete pattern that colocalized with CXCL4, likely in intracellular compartments (Fig. 6a and Extended Data Fig. 11a). Myeloid Differentiation factor 2 (MD-2) associates with TLR4 and is requisite for LPS signaling via TLR4^38, 39^. MD-2 also colocalized with CXCL4 in human macrophages after 30 min stimulation with CXCL4 (Fig. 6b and Extended Data Fig. 11b). Inhibition of MD-2 using MD2-IN-1^40^ abolished both CXCL4 and LPS-induced gene expression (Fig. 6c). These data suggest that CXCL4 and the TLR4/MD-2 complex traffic to similar intracellular compartments after CXCL4 stimulation of human macrophages, where TLR4-TRIF signaling can be activated, and provide an additional two lines of evidence supporting a role for TLR4 in CXCL4 responses.

## Discussion

The importance of CXCL4 in inflammatory disease pathogenesis is well established, especially for inflammation-associated fibrosis in SSc^3, 7, 11, 41, 42, 43^, but the mechanisms by which CXCL4 activates cells are not well defined. Our understanding of how this chemokine activates inflammatory instead of chemotactic responses has been hampered by lack of knowledge of which receptor(s) mediate CXCL4-induced inflammatory responses, and limited characterization of CXCL4-activated signaling pathways. This study advances the field in two ways. First, we have identified that CXCL4 induces a TBK1-JNK signaling axis that complements NF-κB in activating inflammatory, fibrotic and neutrophil chemokine genes; hence, this work highlights a role for TBK1 distinct from its established function in activating IFN responses. CXCL4 also activates STAT3 and RIPK3 that are important in regulating inflammatory responses and cell death. Second, we found that CXCL4 responses were dependent on TLR4, but that CXCL4 induces a gene expression profile distinct from that induced by LPS, suggesting that CXCL4 may modulate TLR4 responses induced by itself or by endotoxin contaminants in the CXCL4 preparations. These findings provide new insights into the functions and mechanisms of action of CXCL4, and suggest a new paradigm by which a chemokine can activate macrophages.

Although a role for TLR4 in monocyte responses to CXCL4 is unexpected, we have provided 6 distinct lines of evidence supporting this notion: 1. Sensitivity of CXCL4 responses to a small molecule TLR4 inhibitor. 2. Inhibition of CXCL4 responses by a TLR4 blocking antibody. 3. Attenuation of CXCL4 responses by siRNA-mediated knockdown of TLR4. 4. Loss of CXCL4-induced gene expression by genetic deletion of *Tlr4*. 5. Attenuation of CXCL4 responses by an MD-2 inhibitor. 6. Colocalization of CXCL4 with TLR4 and MD-2. Collectively, these results make it unlikely that the role of TLR4 is related to off-target or nonspecific effects of the inhibitors or genetic approaches that were used. It is possible that binding of CXCL4 at the plasma membrane by PGs^12^, which increases local CXCL4 concentrations, promotes low affinity interactions with TLR4 and/or MD-2, similar to CXCL4 interactions with the LDLR and CD11b/CD18 previously described^20, 21^. As CXCL4 belongs to the subgroup of chemokines that can self-aggregate^44^, CXCL4 aggregates could crosslink TLR4 and initiate signaling and internalization. In this model, CXCL4 would act as a TLR4 ‘agonist’, as has been proposed for a various endogenous proteins that are DAMPs or alarmins, such as biglycan, hyaluronin, versican, tenascin-C, fibrinogen, heat shock proteins, HMGB1, β-defensins and LDLs^23, 24^. The possibility that CXCL4 is a direct TLR4 agonist would require extensive validation in future work using biochemical and structural biology approaches with purified proteins.

Another non-mutually exclusive and equally interesting possibility about how TLR4 is activated in our system is based on observations that many of the endogenous proteins suggested to activate TLR4, including heat shock proteins, surfactant protein A, β-defensins, fibrinogen, biglycan and LDLs can bind and/or enhance the sensitivity of cells to LPS in a manner analogous to CD14^23^. In this model, these alarmins serve as enhancers of LPS sensing, rather than direct agonists of TLR4. In our system, addition of CXCL4 to cell cultures resulted in only extremely low LPS concentrations (1-2 pg/ml), which is not sufficient to activate monocytes when LPS alone added to such cultures^12^. However, testing of additional lots of CXCL4 from various vendors showed variation in endotoxin contamination, including higher concentrations that could contribute to monocyte activation. Although LPS-binding by proteins can suppress the readout of the LAL assay, the complete loss of bioactivity after heating CXCL4 solutions at 100 °C for 10 minutes to denature CXCL4 supports the importance of CXCL4 itself in eliciting the monocyte response. Although heat denaturation of LPS typically requires dry heat (‘burning’) at >250 °C for > 30 min, we found that heating could affect the activity of low concentrations of LPS. Despite extensive additional efforts to address endotoxin contamination, we were not able to completely dissect the contributions of CXCL4 alone and endotoxin contamination to the observed results. Notably, the CXCL4-induced signaling and gene induction events observed herein in monocytes were not observed in pDCs or B cells^18, 45^, which express minimal TLR4 levels and are much less sensitive to TLR4 ligands.

The transcriptional output after CXCL4 stimulation differed from the transcriptional output of TLR4 activation by LPS, most notably a negligible IFN response and lack of activation of IL-12 family genes by CXCL4. A difference in macrophage signaling and transcriptional responses to an endogenous putative TLR4 agonist and LPS has been previously described for tenascin-C^46^. Similar to CXCL4, tenascin-C was not able to activate IL-12 family genes; IFN responses were not analyzed in this previous study. As TLR responses and the balance between inflammatory and IFN responses are affected by intracellular trafficking through endolysosomal compartments^22, 47^, it is possible that altered trafficking of TLR4 after CXCL4 stimulation, as suggested by Figure 7, affected the IFN response by modulating the intensity or duration of IRF activation and ‘repurposing’ TBK1 for inflammatory gene induction. Alternatively, signaling crosstalk with the additional receptors engaged by CXCL4, such as CD11b/CD18, LDLRs and cell surface PGs^12, 17, 20, 21^, could affect TLR4-mediated transcriptional output. CD11b/CD18 is well established to modulate TLR4 signaling in a complex manner^48, 49^. Another consideration is that the differences may reflect a cell-intrinsic difference in responses to strong versus weak stimulation of TLR4; in the latter scenario it is difficult to exclude a contribution from contaminating endotoxin.

CXCL4, TLR8 and TLR4 have been shown to play important roles in tissue fibrosis in diseases such as SSc^3, 11, 43, 50, 51, 52^ and can affect the activation and function of various pathogenic cell types. One pathogenic function of CXCL4 in SSc is to promote NA-mediated activation of TLR8 and TLR9 in plasmacytoid DCs^3, 18, 52^. A key role for TLR4 in SSc pathogenesis has been established by the efficacy of TLR4 inhibitors or genetic deletion of *Tlr4* in ameliorating disease in preclinical models^50, 51^. In contrast to NA-mediated activation of TLR8, in SSc TLR4 is activated by endogenous protein ‘agonists’ including tenacin-C, fibronectin-EDA, HMGB1 and S100 proteins. Our findings suggest that the high concentrations of CXCL4 present in SSc may also contribute to pathogenesis by activation, either directly or indirectly, of TLR4.

In summary, CXCL4 activation of human monocytes diminished when TLR4 was blocked and involved NF-κB and TBK1-JNK signaling pathways that drive inflammatory and tissue repair/fibrotic gene expression. CXCL4 also activates expression of neutrophil chemokine genes. These findings provide insights about how CXCL4 can contribute to inflammatory and fibrotic disease pathogenesis, and suggest strategies for targeting CXCL4-activated pathways to suppress pathology.

## Methods

### Human cells

Deidentified human buffy coats were purchased from the New York Blood Center following a protocol approved by the Hospital for Special Surgery Institutional Review Board. Peripheral blood mononuclear cells (PBMCs) were isolated using density gradient centrifugation with Lymphoprep (Accurate Chemical, Carle Place, NY, USA) and monocytes were purified with anti-CD14 magnetic beads from PBMCs immediately after isolation as recommended by the manufacturer (Miltenyi Biotec)^53^. Monocytes were cultured overnight at 37°C, 5% CO_2_ in RPMI-1640 medium (Invitrogen) supplemented with 10% heat-inactivated defined FBS (HyClone Fisher), penicillin-streptomycin (Invitrogen), L-glutamine (Invitrogen) and 20 ng/ml human M-CSF. Then, the cells were treated as described in the figure legends.

### Mouse bone marrow-derived macrophage (BMDM) culture

Bone marrow cells from male C57BL/6J mice and male *Tlr4^-/-^* mice were harvested after euthanasia by CO2 asphyxiation, and cultured in RPMI-1640 medium (Invitrogen) supplemented with 10% heat-inactivated defined FBS (HyClone Fisher), penicillin-streptomycin (Invitrogen), L-glutamine (Invitrogen) and 20 ng/ml mouse M-CSF 6 days. Then, the cells were treated as described in the figure legends.

### RNA sequencing

Libraries for sequencing were prepared using extracted RNA and the NEBNext Ultra II RNA Library Prep Kit for Illumina following the manufacturer’s instructions (New England Biolabs). Quality of all RNA and library preparations were evaluated with BioAnalyzer 2100 (Agilent).

Human RNA libraries were sequenced by the Genomics Resources Core Facility at Weill Cornell using a HiSeq4000, 50-bp single-end reads at a depth of –at least 20 million uniquely mapped reads per sample. Read quality was assessed and adapters trimmed using Fastp. Reads were then mapped to the human genome (hg38) and mouse genome (mm10) and reads in exons were counted against Gencode M25 with STAR Aligner. Differential gene expression analysis was performed in R using edgeR. Genes with low expression levels (< 3 counts per million in at least one group) were filtered from all downstream analyses. Benjamini-Hochberg false discovery rate (FDR) procedure was used to correct for multiple testing.

### Analysis of mRNA amounts

Total RNA was extracted with a RNeasy Mini Kit (QIAGEN) and was reverse-transcribed with a RevertAid RT Reverse Transcription Kit (Thermo Fisher Scientific, Catalog number: K1691).

Real-time PCR was performed with Fast SYBR Green Master Mix and a 7500 Fast Real-time PCR system (Applied Biosystems). The primer sequences for the qPCR reactions are listed in Extended Data Table 1. CT values of target genes were normalized to *GAPDH* expression and are shown as percentage of GAPDH (100/2^^ΔCt^).

### Western Blotting

Cells were lysed in 50 µl of cold buffer containing 50 mM Tris-HCl pH 7.4, 150 mM NaCl, 1 mM EDTA, 1% (vol/vol) Triton X-100, 2 mM Na3VO4, 1x phosSTOP EASYPACK, 1 mM Pefabloc, and 1× EDTA-free complete protease inhibitor cocktail (Roche, Basel, Switzerland), and incubated for 10 min on ice. Then, cell debris was pelleted at 13,000 rpm at 4°C for 10 min. The soluble protein fraction was mixed with 4× Laemmli Sample buffer (BIO-RAD, Cat. #1610747) and 2-mercroptoehanol (BME) (Sigma-Aldrich). Samples for Western blotting were subjected to electrophoresis on 4–12% Bis-Tris gels (Invitrogen). Proteins were transferred to polyvinylidene difluoride membrane was as previously reported^54^. Membranes were blocked in 5% (w/v) Bovine Serum Albumin in TBS (20 mm Tris, 50 mm NaCl, pH 8.0) with 0.1% (v/v) Tween 20 (TBST) at room temperature for at least 1 h with shaking at 60 rpm. Membranes were then incubated with primary antibodies at 4 °C overnight with shaking at 60 rpm. Membranes were washed 3 times in TBST, then probed with anti-mouse or anti-rabbit IgG secondary antibodies conjugated to horseradish peroxidase (GE Healthcare, cat: NA9310V and NA9340V) diluted in TBST at room temperature for one hour with shaking at 60 rpm. Next, membranes were washed 3 times in TBST at room temperature for with shaking at 60 rpm. Antibody binding was detected using enhanced chemiluminescent substrates for horseradish peroxidase (HRP) (ECL Western blotting reagents (PerkinElmer, cat: NEL105001EA) or SuperSignal West Femto Maximum Sensitivity Substrate (Thermo Fisher Scientific, cat: 34095), according to the manufacturer’s instructions, and visualized using premium autoradiography film (Thomas Scientific, cat: E3018). To detect multiple proteins on the same experimental filter, membranes were cut horizontally based on the molecular size of the target proteins. For membranes that required probing twice or more using different primary antibodies, RestoreTM PLUS Western blotting stripping buffer (Thermo Fisher Scientific) was applied on the blots with shaking at 60 rpm for 15 min following previous development. Antibodies used are identified in Extended Data Table 2.

### Immunoprecipitation

Human monocytes (20 x 10^6^) were lysed in 650 µl of IP Lysis/Wash buffer (Pierce Direct IP Kit, Thermo Fisher Scientific, Cat. #26148) and the supernatants of cell lysates after centrifugation were transferred to columns containing the TBK1 antibody linked to AminoLink Plus Coupling Resin to pull down the proteins interacting with TBK1 using the Pierce Direct IP Kit according to the manufacturer’s instructions. The immunoprecipitated proteins were denatured with 4× Laemmli Sample buffer and BME and boiled for 5 min, resolved by SDS-PAGE and immunoblotted with indicated antibodies.

### RNA interference

For RNA interference (RNAi) experiments, primary human monocytes (6 x 10^6^ cells) were transfected with 0.1 nmol of siRNA oligonucleotides (listed in Extended Data Table 3) using a Human Monocyte Nucleofactor Kit (Lonza, VVPA-1007) and the AMAXA Nucleofector System (Lonza) program Y001 for human monocyte transfection according to the manufacturer’s instructions.

### Endotoxin detection

The concentration of endotoxin in CXCL4 stock solutions was measured by Chromogenic LAL Endotoxin Assay Kit (GenScript, cat. No: L00350C) according to the manufacturer’s instructions.

### Confocal microscopy

The protocol was as described in our previous publication^55^. Briefly, human monocytes were cultured in 20 ng/ml M-CSF for 3 to 6 days and incubated with 10 μg/ml human CXCL4 for 30 min, fixed in PBS buffer containing 4% paraformaldehyde at room temperature (RT) for 15 min, and permeabilized for 10 min with 0.1 % Triton-X 100 in PBS. Samples were then blocked by 3% BSA in PBS for 30 min at RT, and labeled with anti-CXCL4-PE (1:50) (R&D system, IC7952P) and anti-TLR4 (10 µg/ml) (R&D system, AF1478), or anti-MD-2 (2 µg/ml) (Thermo Fisher Scientific, PA5-20058) for 3 h at RT. Secondary antibodies (1:1000) anti-goat-AF647 (1:1000) (Thermo Fisher Scientific, A-21446) or anti-rabbit CF647 (1:1000) (Sigma, SAB4600175) are used when needed. Images were acquired using a ZEISS LSM880 confocal microscope and a 63×/1.4 N.A. objective.

### Statistical Analysis

Graphpad Prism for Windows was used for all statistical analysis. Information about the specific tests used, and number of independent experiments is provided in the figure legends. Two-way ANOVA with Sidak correction for multiple comparisons was used for grouped data; when the data did not pass normality distribution by the Shapiro-Wilk test, the Friedman test with Dunn’s correction was used. Otherwise, one-way ANOVA with the Geisser-Greenhouse correction and Tukey’s post hoc test for multiple comparisons was performed. For paired data, when the data did not pass the normal distribution by F test the Wilcoxon signed-rank test was performed, otherwise, paired t test was used.

### Data availability

The RNAseq data sets that support the findings of this study and were generated by the authors as part of this study have been deposited in the Gene Expression Omnibus database with the accession code: GSE210105. Otherwise, the data is either contained within the manuscript or available from the authors on request.

## Acknowledgments

We thank the Weill Cornell Medicine Genomics Core Facility for next generation sequencing, the Weill Cornell Medicine – HSS Flow Cytometry Core Facility for flow cytometry support, and David Oliver (HSS Genomics Center) for advice and discussions. This work was supported by NIH grants R01 AI044938, R01 AR46713 and R01 AR050401 (L.B.I.). F.J.B. is supported by grants from the NIH 1R01AI132447, from the Scleroderma Research Foundation and from the Scleroderma Foundation. The David Z. Rosensweig Genomics Center at HSS is supported by The Tow Foundation. All NIH-supported work was performed at HSS.

## Author contributions

C.Y. conceptualized, designed, and performed most of the experiments, performed bioinformatic analysis, prepared figures and wrote the manuscript. R.Y., B. M., R. B., Y. Z., Y. D., M.D.A.K., contributed experiments or experimental expertise; F.J.B. contributed expertise and intellectual input and edited the manuscript. L.B.I. conceptualized and oversaw the study and edited the manuscript. All authors reviewed and provided input on the manuscript.

