## Supplemental materials for "CXCL4 signaling and gene induction in human monocytes"

**Extended Data Figures: 11**

**Extended Data Table: 2**

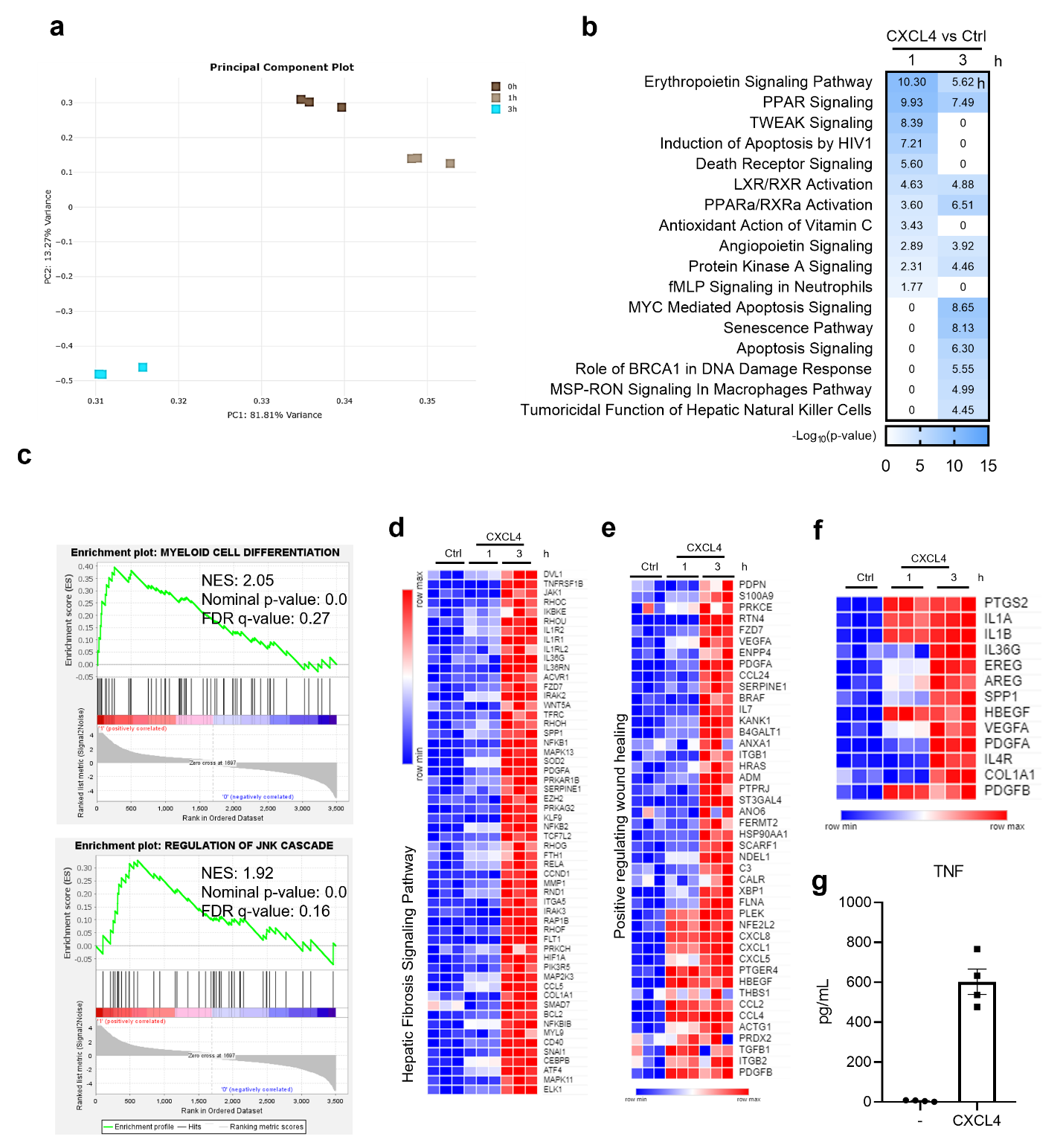

**Extended Data Figure 1.** (a) Principal Component Analysis (PCA) of RNAseq data. (b) IPA analysis showing significant enrichment of suppressed pathways regulated by CXCL4 1 h or 3 h post treatment. (c) The GSEA gene set enrichment analyis using the DEGs of the CXCL3 h treatment. Heatmaps of Hepatic Fibrosis Signaling Pathway (d), Positive regulating wound healing (e) and fibrosis-related genes (f). (g) ELISA to measure cytokine TNF protein secretion in the supernatant of human monocytes stimulated with CXCL4 for 6 h (n = 4 independent donors). Data are depicted as mean ± SEM. Source data are provided as a Source Data file.

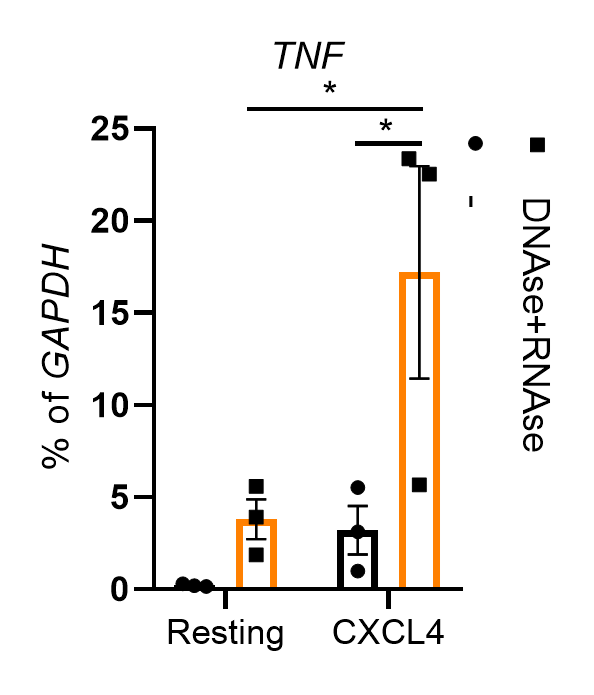

**Extended Data Figure 2.** Human primary monocytes were isolated with CD14^+^ magnetic beads and rested overnight with M-CSF (20 ng/ml). mRNA of *TNF* measured by qPCR and normalized relative to *GAPDH* mRNA in cells pre-treated with DNase I (1000 U) + RNase A (50 ug/ml) for 1 h and then stimulated with CXCL4 for 3 h (n = 3 independent donors). Data are depicted as mean ± SEM. **p ≤ 0.01; *p ≤ 0.05 by Two-way ANOVA. Source data are provided as a Source Data file.

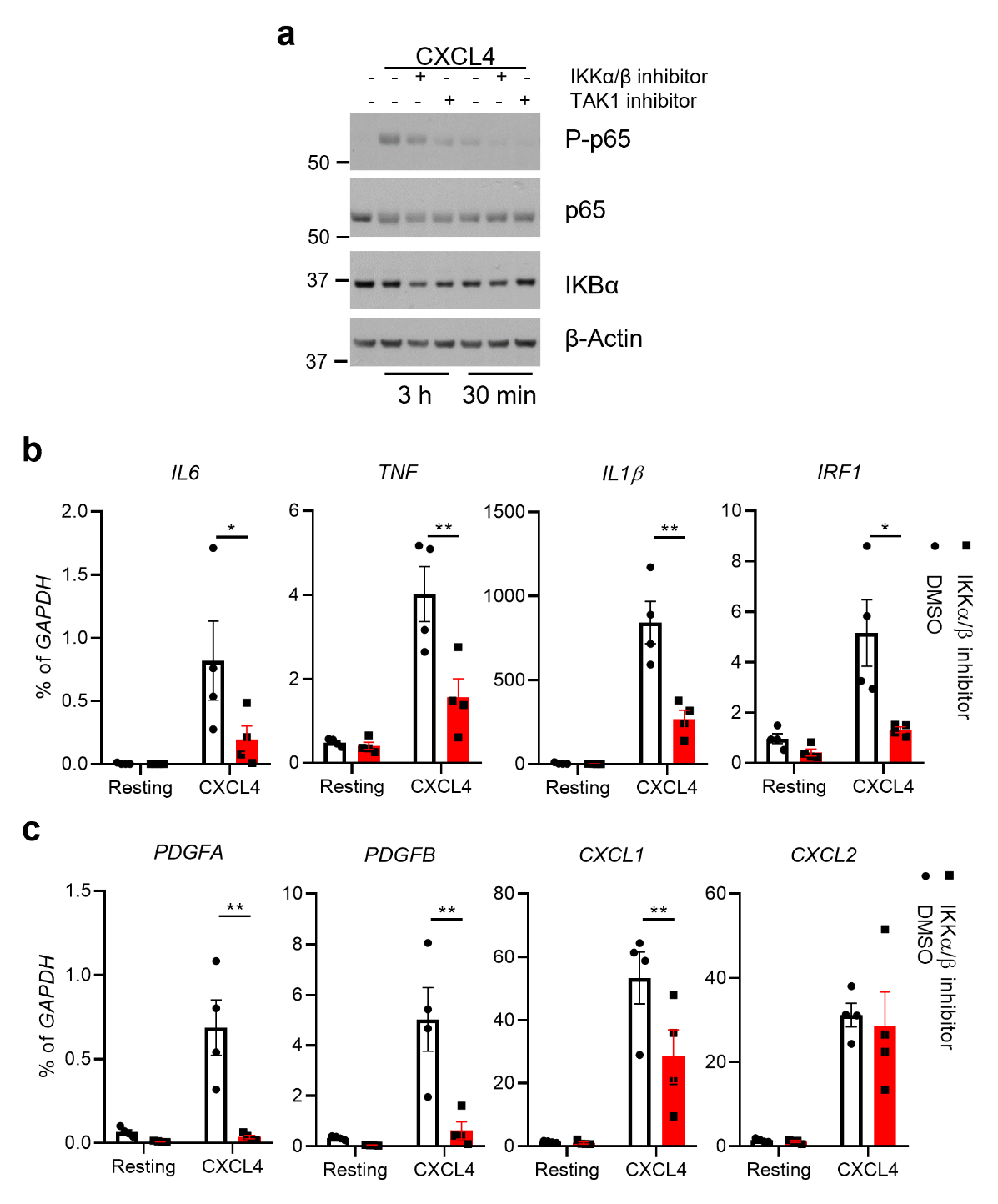

**Extended Data Figure 3.** (a) Immunoblot of phospho-p65, p65, phospho-IKBα and total IKBα with whole cell lysates of human primary monocytes stimulated with CXCL4 in the presence/absence of either BMS-345541 or TAK1 inhibitor Takinib (10 µM) for 3 h (representative of 3 independent donors). mRNA of inflammatory genes (b) and tissue repair/fibrotic genes (c) measured by qPCR and normalized relative to *GAPDH* mRNA in human primary monocytes pre-treated with IKKα/β inhibitor BMS-345541 (10 µM) for 30 min and then stimulated with CXCL4 for 3 h (n = 4 independent donors). Data are depicted as mean ± SEM (b, c). **p ≤ 0.01; *p ≤ 0.05 by Two-way ANOVA. Source data are provided as a Source Data file.

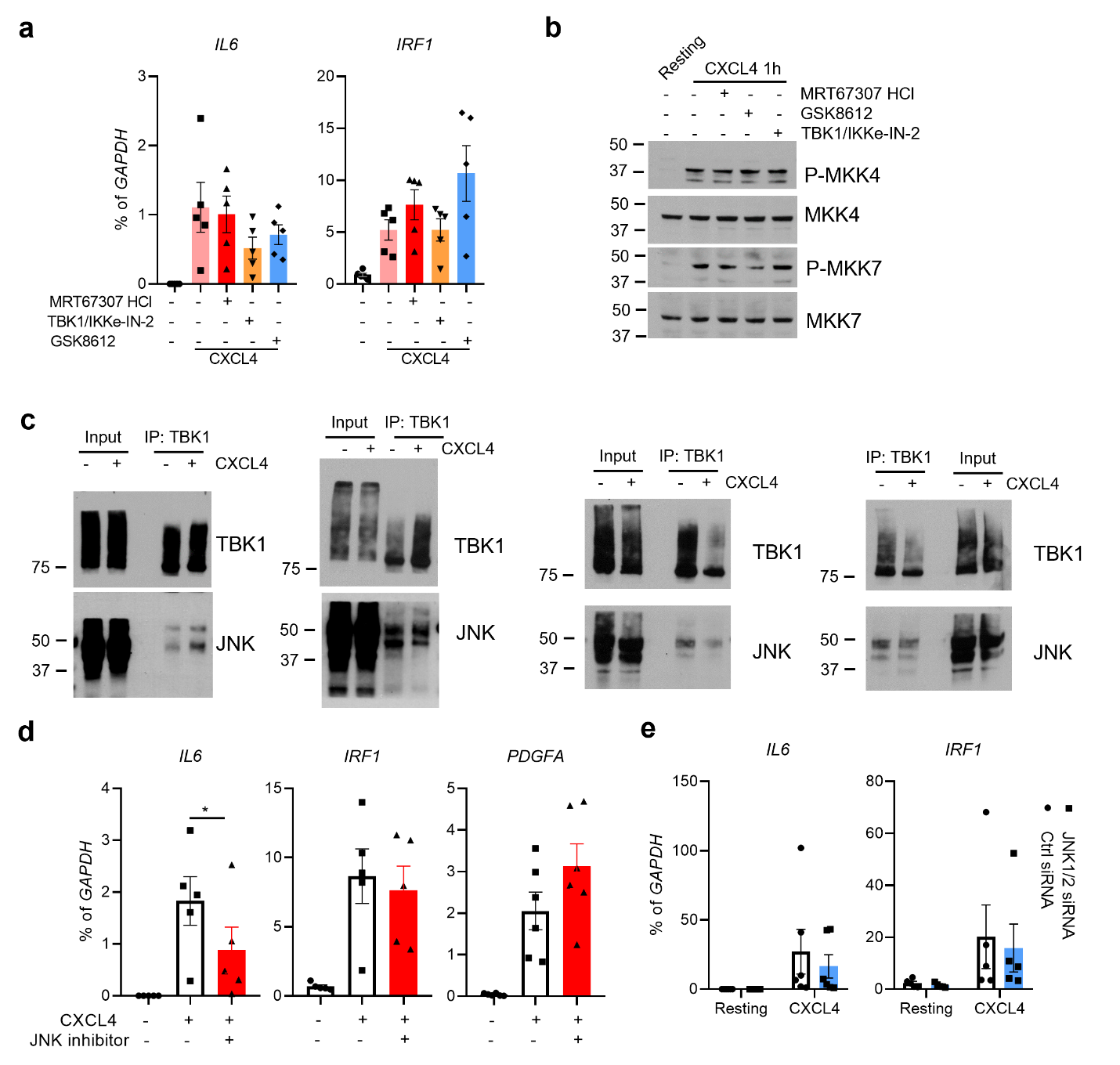

**Extended Data Figure 4.** (a) mRNA of target genes measured by qPCR and normalized relative to *GAPDH* mRNA in human primary monocytes pre-treated with TBK1/IKKε inhibitors MRT67307 HCl (10 µM), TBK1/IKKε-IN-2 (1 µM) or GSK8612 (50 µM) for 30 min and then stimulated with CXCL4 for 3 h (n = 5 independent donors). (b) Immunoblot of phospho-MKK4/7 and MKK4/7 with whole cell lysates of human primary monocytes stimulated with CXCL4 in the presence/absence of MRT67307 HCl (10 µM), TBK1/IKKε-IN-2 (1 µM) or GSK8612 (50 µM) for 1 h (n = 2 independent donors). (c) Immunoblot of TBK1 and JNK from whole cell lysates immunoprecipitated with TBK1 antibodies (n = 5 independent donors; the results with 4 donors are shown here and the 5^th^ donor on Fig. 2d). (d, e) mRNA of indicated genes measured by qPCR and normalized relative to *GAPDH* mRNA in human primary monocytes pre-treated with JNK inhibitor SP600125 (10 µM) (d) for 30 min or 3 days after nucleofection of monocytes with control or JNK1 + 2-specific siRNAs (e) and then stimulated with CXCL4 for 3 h (n = 5 independent donors for e). Data are depicted as mean ± SEM (a, d, e). *p ≤ 0.05 by One-way ANOVA (d). Source data are provided as a Source Data file.

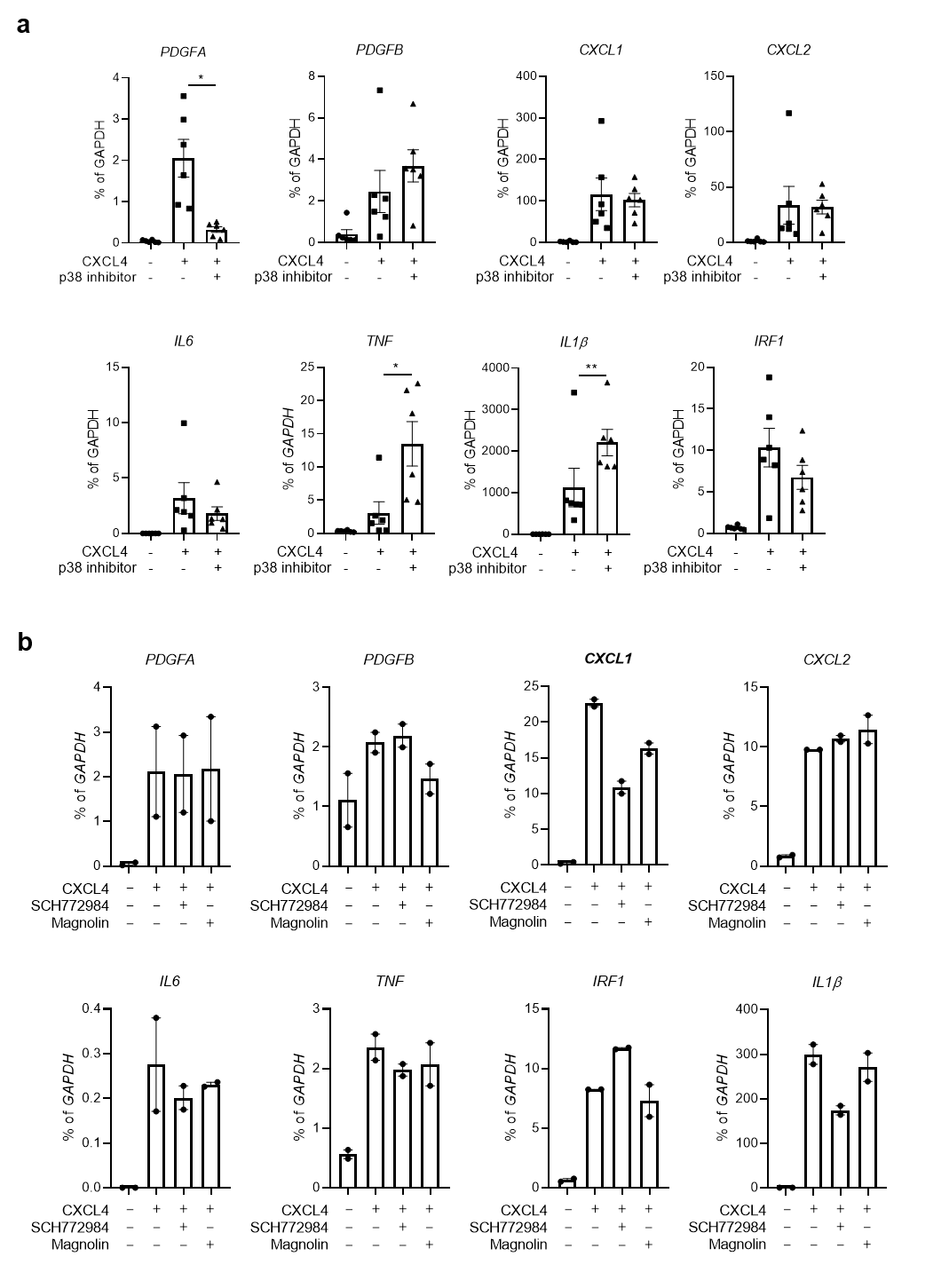

**Extended Data Figure 5.** (a) mRNA of indicated genes measured by qPCR and normalized relative to *GAPDH* mRNA in human primary monocytes pre-treated with p38 inhibitor SB 202190 (10 µM) for 30 min and then stimulated with CXCL4 for 3 h (n = 6 independent donors). (b) mRNA of indicated genes measured by qPCR and normalized relative to *GAPDH* mRNA in human primary monocytes pre-treated with ERK inhibitors SCH772984 (10 µM) and Magnolin (10 µM) for 30 min and then stimulated with CXCL4 for 3 h (n = 2 independent donors). Data are depicted as mean ± SEM. **p ≤ 0.01; *p ≤ 0.05 by One-way ANOVA (a). Source data are provided as a Source Data file.

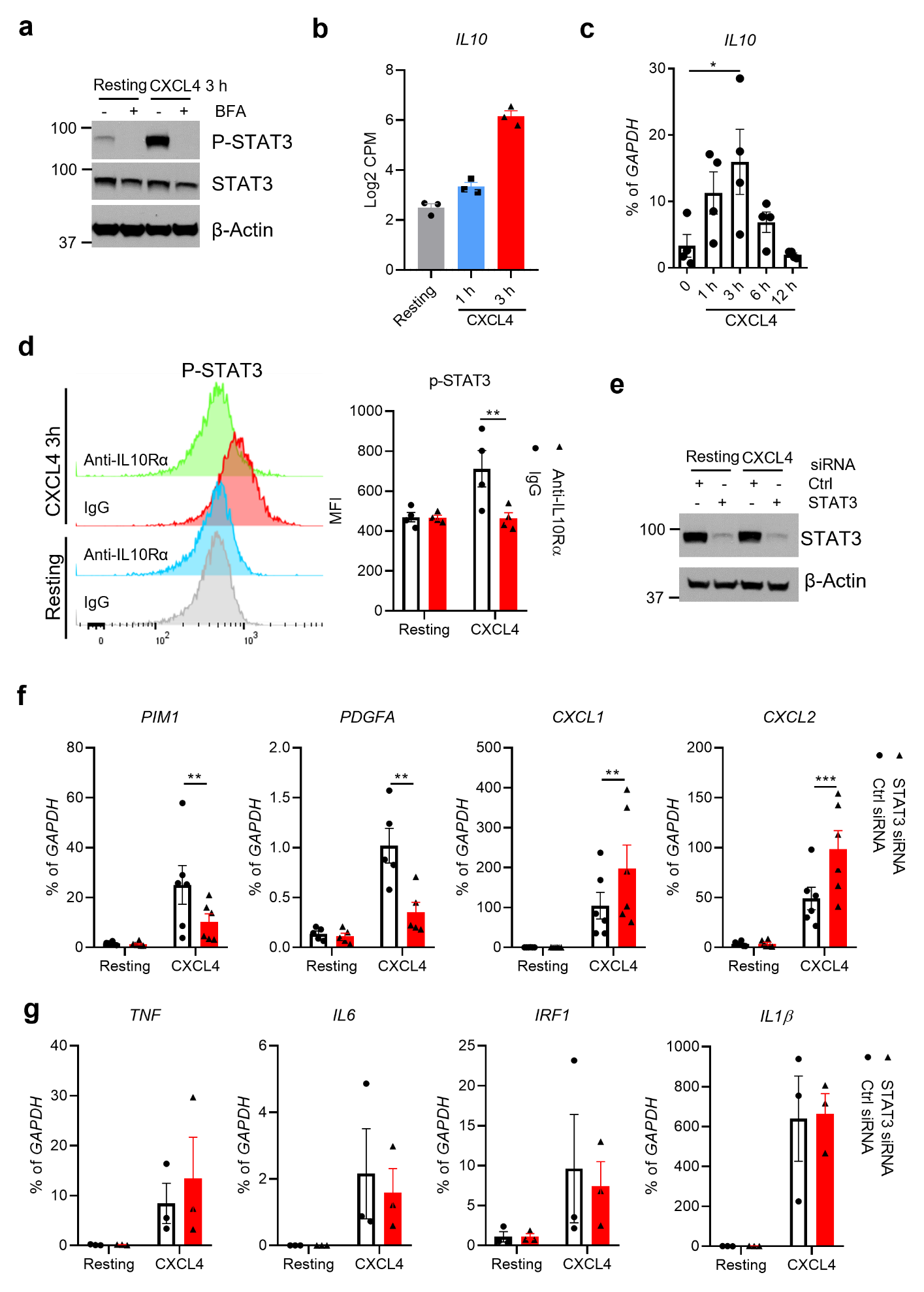

**Extended Data Figure 6.** (a) Immunoblot of phospho-STAT3, STAT3 with whole cell lysates of human primary monocytes stimulated with CXCL4 in the presence/absence of BFA for 3 h (representative of 2 independent donors). (b) Log2 CPM of *IL10* expression from RNA-seq. (c) mRNA of *IL10* measured by qPCR and normalized relative to *GAPDH* mRNA in human primary monocytes stimulated with CXCL4 in time-course experiments (n = 4 independent donors). (d) Flow cytometric analysis of STAT3 phosphorylation in cells stimulated with CXCL4 for 3 h in the absence or presence of IL10Rα neutralizing antibody (10 µg/ml). Left panel, representative FACS plot; right panel, cumulative data (n = 4 independent donors). (e) Immunoblot of STAT3 using whole cell lysates 3 days after nucleofection of monocytes with control or STAT3-specific siRNAs (representative of 3 independent donors). (f) mRNA of indicated genes was measured by qPCR and normalized relative to *GAPDH* mRNA after knockdown of STAT3 by siRNA for 3 days (n = 6 independent donors). (g) mRNA of indicated genes was measured by qPCR and normalized relative to *GAPDH* mRNA after knockdown of STAT3 by siRNA for 3 days (n = 3 independent donors). Data are depicted as mean ± SEM (b, c, d, f, g). ***p ≤ 0.001; **p ≤ 0.01; *p ≤ 0.05 by One-way ANOVA (c) or Two-way ANOVA (d, f). Source data are provided as a Source Data file.

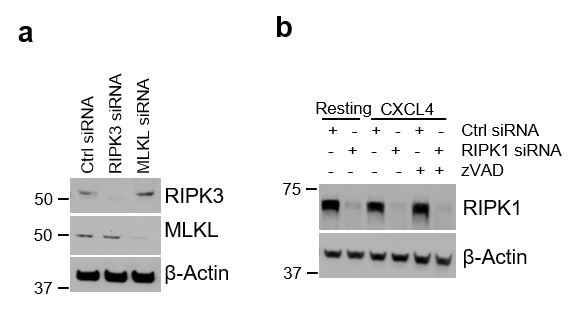

**Extended Data Figure 7.** (a, b) Immunoblots of cells 3 days after nucleofection of monocytes with control (Ctrl) siRNA, RIPK3 and MLKL (a) or RIPK1 (b) specific siRNAs in the indicated conditions (representative of 3 independent donors). Source data are provided as a Source Data file.

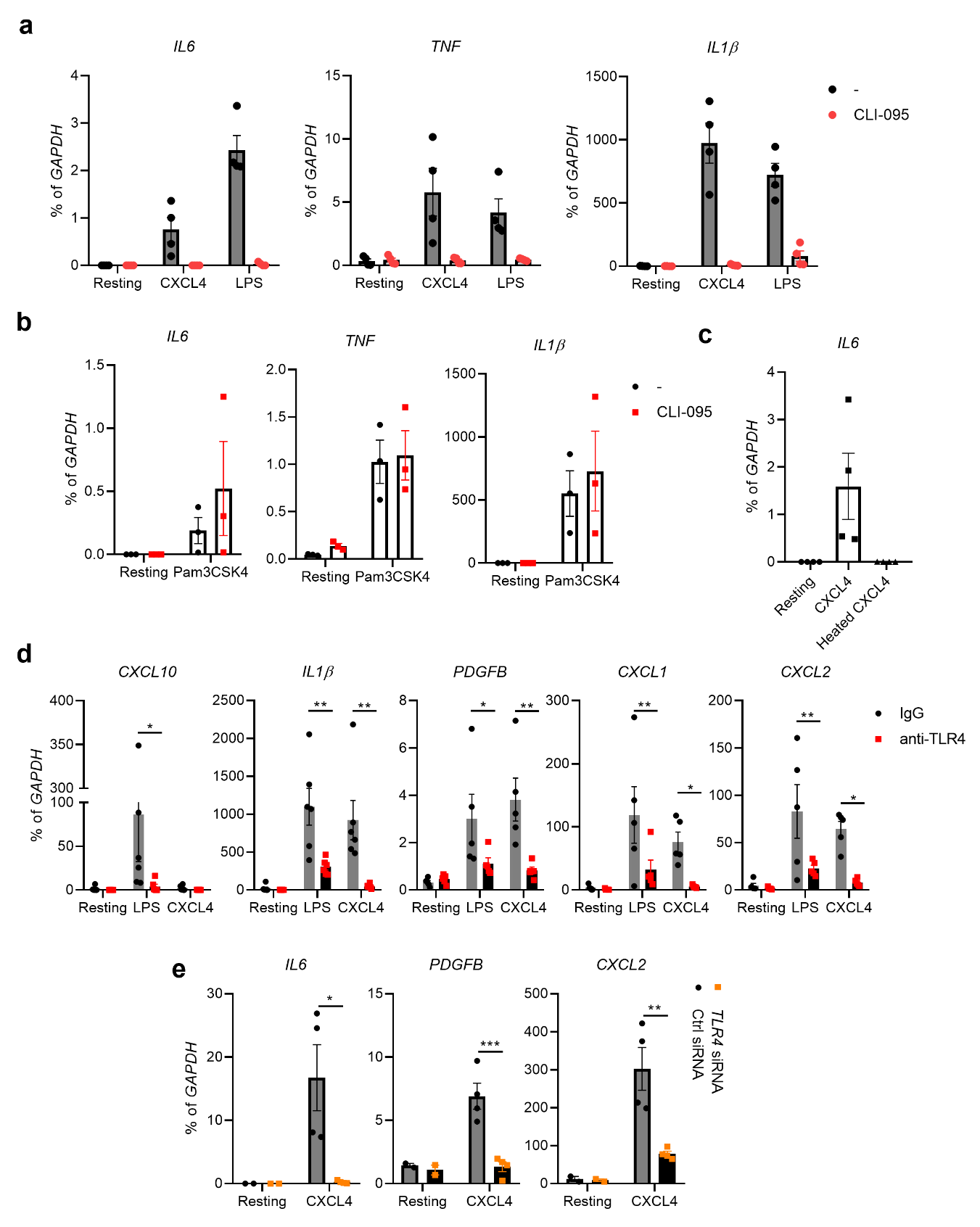

**Extended Data Figure 8.** (a, b) mRNA of *IL6*, *TNF* and *IL1β* measured by qPCR and normalized relative to GAPDH mRNA in human primary monocytes pre-treated with TLR4 inhibitor CLI-095 (2.5 µM) for 1 h and then stimulated with CXCL4 or LPS (5 ng/ml) (a) or Pam3CSK4 (20 ng/ml) for 3 h (n = 4 independent donors for a, 3 independent donors for b). (c) mRNA of *IL6* measured by qPCR and normalized relative to *GAPDH* mRNA in human primary monocytes treated with CXCL4 or heat denatured CXCL4 for 3 h (n = 4 independent donors). (d) mRNA of indicated genes measured by qPCR and normalized relative to *GAPDH* mRNA in human primary monocytes pre-treated with TLR4 neutralizing antibody (5 µg/ml) for 1 h and then stimulated with CXCL4 or LPS (5 ng/ml) for 3 h (n = 6 donors for *CXCL10* and *IL1β*, 5 independent donors for the rest). (e) mRNA of indicated genes was measured by qPCR and normalized relative to *GAPDH* mRNA in human primary monocytes after knockdown of TLR4 by specific siRNA for 3 days and then stimulated with CXCL4 for 3 h (n = 4 independent donors). Data are depicted as mean ± SEM (a - e). ***p ≤ 0.001; **p ≤ 0.01; *p ≤ 0.05 by Two-way ANOVA (d, e). Source data are provided as a Source Data file.

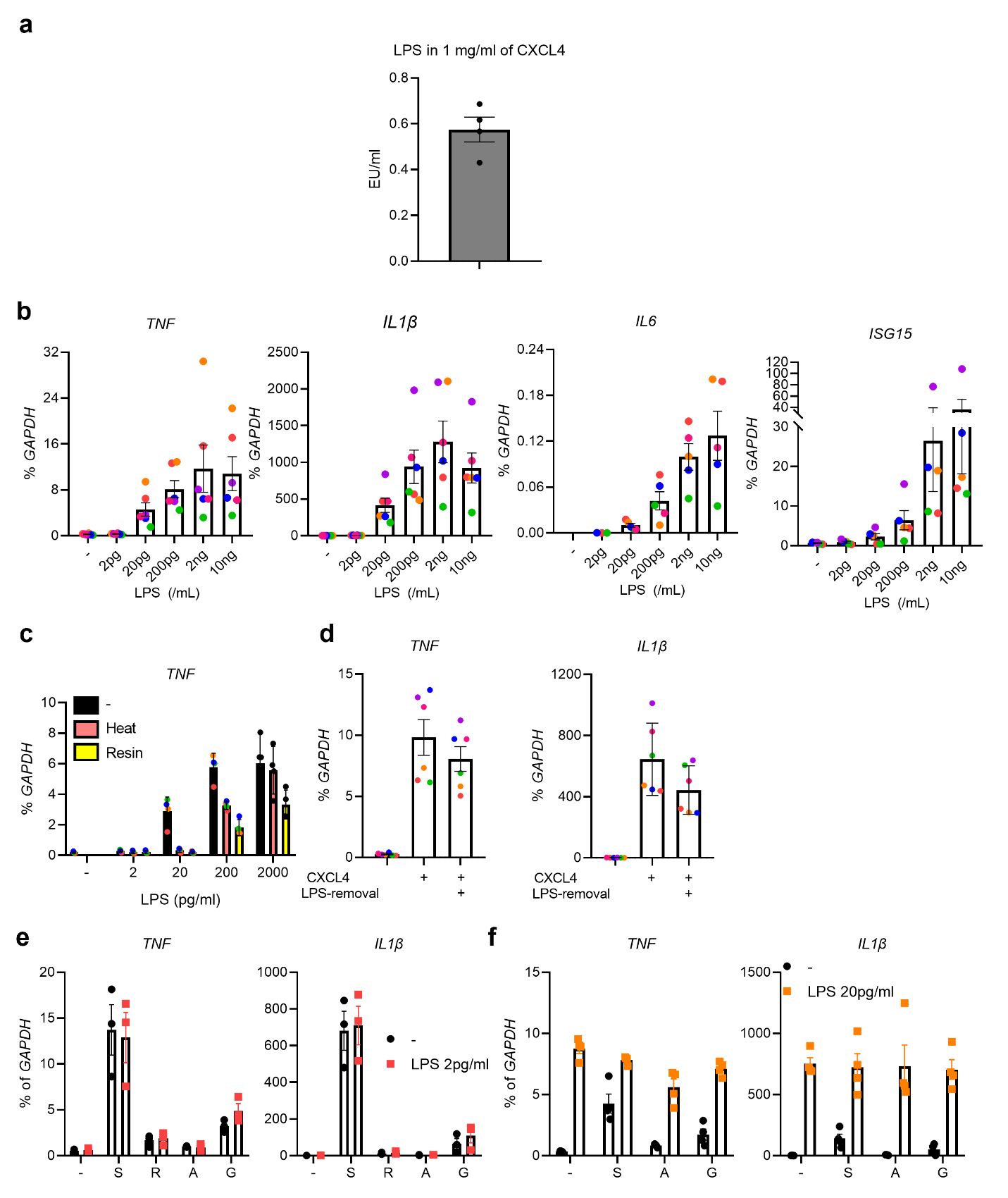

**Extended Data Figure 9.** Endotoxin assay of CXCL4 stock solution (n = 4, 2 technical replicates from 2 batches of CXCL4). Source data are provided as a Source Data file.

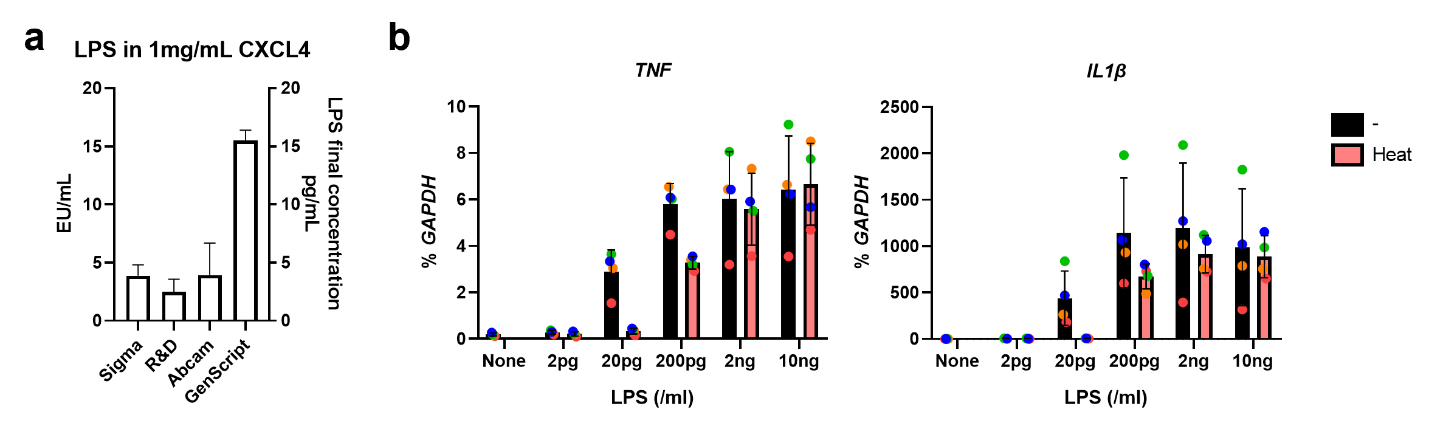

**Extended Data Figure 10.** (a) Endotoxin levels were measured by LAL assay in 1 mg/ml CXCL4 from different sources. (b) mRNA of indicated genes measured by qPCR and normalized relative to *GAPDH* mRNA in human primary monocytes treated with LPS or heated LPS in the indicated concentration (n = 4 independent donors). Data are depicted as mean ± SEM. Source data are provided as a Source Data file.

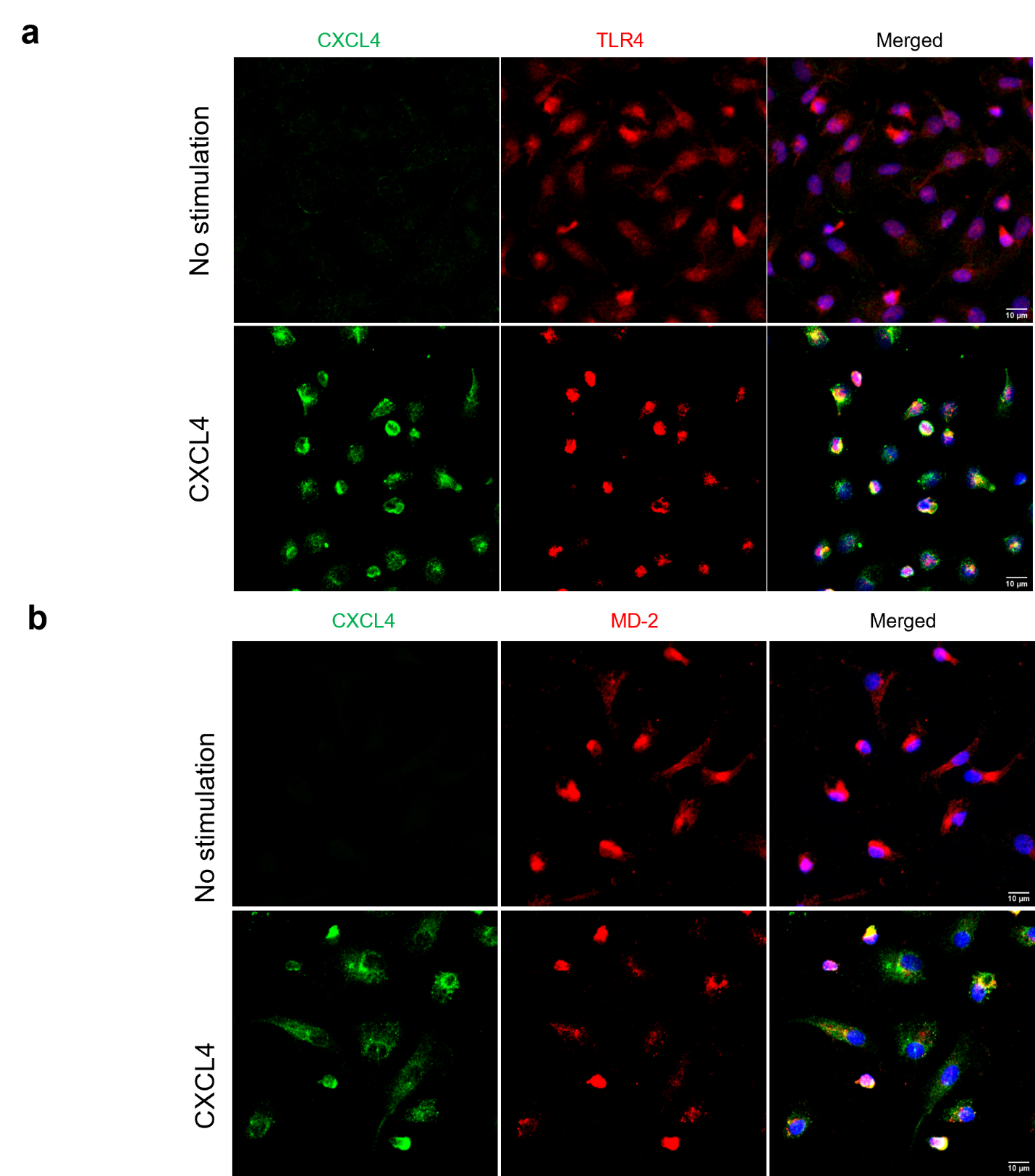

**Extended Data Figure 11.** (a, b) Representative confocal images of CXCL4 colocalization with TLR4 (a) and MD-2 (b) in human primary monocytes 30 min after treatment (n = 3 independent donors for each experiment).

**Extended Data Table 1: Primers and siRNAs.**

| **Primers for qPCR Source** | | |
| --- | --- | --- |
| Human *GAPDH* F: ATCAAGAAGGTGGTGAAGCA; R: GTCGCTGTTGAAGTCAGAGGA | This paper |  |
| Human *IL6* F: TAATGGGCATTCCTTCTTCT; R: TGTCCTAACGCTCATACTTTT | This paper |  |
| Human *TNF* F: AATAGGCTGTTCCCATGTAGC; R: AGAGGCTCAGCAATGAGTGA | This paper |  |
| Human *IL1β* F: TTCGACACATGGGATAACGAGG; R: TTTTTGCTGTGAGTCCCGGAG | This paper |  |
| Human *IL12A* F: GATGGCCCTGTGCCTTAGTA; R: TCAAGGGAGGATTTTTGTGG | This paper |  |
| Human *IL12B* F: AAGCACTTTGGAGGAAGCATAG; R: TGAACCCAAGTCAATGTGAGTC | This paper |  |
| Human *CXCL10* F: TTAATCTTGTCTCTGGGCTTGG; R: GTTGGGGAATGAGGTTAGGG | This paper |  |
| Human *IRF1* F: GCACTAAGCGAAAATTGCA; R: GGGAGTTTCCTTCACATTCA | This paper |  |
| Human *PIM1* F: GGCTCGGTCTACTCAGGCA; R: GGAAATCCGGTCCTTCTCCAC | This paper |  |
| Human *ISG15* F: TGGACAAATGCGACGAACCTC; R: TCAGCCGTACCTCGTAGGTG | This paper |  |
| Human *IFIT1* F: GCGCTGGGTATGCGATCTC; R: CAGCCTGCCTTAGGGGAAG | This paper |  |
| Human *PDGFA* F: GGACAGTGCGACGGTATTT; R: GGAGAAACACAAAGCCAGAAAC | This paper |  |
| Human *PDGFB* F: CTGTCTCTCTGCTGCTACCTG; R: TCGAGTGGTCACTCAGCATC | This paper |  |
| Human *CXCL1* F: AGTCATAGCCACACTCAAGAATGG; R: GATGCAGGATTGAGGCAAGC | This paper |  |
| Human *CXCL2* F: GCATCGCCCATGGTTAAGA; R: TCAGGAACAGCCACCAATAAG | This paper |  |
| Human *IL10* F: TTAATCTTGTCTCTGGGCTTGG; R: GTTGGGGAATGAGGTTAGGG | This paper |  |
| Mouse *Gapdh* F: ATCAAGAAGGTGGTGAAGCA; R: AGACAACCTGGTCCTCAGTGT | This paper |  |
| Mouse *Il6* F: TGGCTAAGGACCAAGACCATCCAA; R: AACGCACTAGGTTTGCCGAGTAGA | This paper |  |
| Mouse *Tnf* F: CCCTCACACTCAGATCATCTTCT; R: GCTACGACGTGGGCTACAG | This paper |  |
| Mouse *Il1β* F: AGCTTCCTTGTGCAAGTGTCT; R: GACAGCCCAGGTCAAAGGTT | This paper |  |
| **Oligonucleotides Source CAS#** | | |
| siGENOME Human MAPK8 siRNA | Horizon | M-003514-04-0005 |
| siGENOME Human MAPK9 siRNA | Horizon | M-003505-02-0005 |
| siGENOME Human STAT3 siRNA | Horizon | M-003544-02-0005 |
| siGENOME Human TLR4 siRNA | Horizon | M-008088-01-0005 |
| ON-TARGETplus Human RIPK1 siRNA | Dharmacon | L-004445-00-0005 |
| SMARTpool: siGENOME RIPK3 siRNA | Dharmacon | M-003534-01-0005 |
| SMARTpool: siGENOME MLKL siRNA | Dharmacon | M-005326-00-0005 |
| siGENOME Human TICAM1 siRNA | Dharmacon | M-012833-02-0005 |

**Extended Data Table 2: Antibodies and Others.**

| **Antibodies Source CAS#** | | |
| --- | --- | --- |
| Stat3 (D3Z2G) (1:1000) | Cell Signaling Technology | 12640S |
| Phospho-Stat3 (Tyr705) (D3A7) XP® Rabbit mAb (PE Conjugate) (1:50) | Cell Signaling Technology | 8119S |
| Phospho-SEK1/MKK4 (Ser257/Thr261) (1:500) | Cell Signaling Technology | 9156S |
| SEK1/MKK4 Antibody (1:1000) | Cell Signaling Technology | 9152S |
| Phospho-MKK7 (Ser271/Thr275) (1:500) | Cell Signaling Technology | 4171S |
| MKK7 Antibody (1:1000) | Cell Signaling Technology | 4172S |
| TBK1/NAK (E8I3G) (1:1000) | Cell Signaling Technology | 38066S |
| Phospho-TBK1/NAK (Ser172) (D52C2) (1:1000) | Cell Signaling Technology | 5483T |
| Phospho-IKKε (Ser172) (D1B7) (1:500) | Cell Signaling Technology | 8766S |
| IKKε (1:500) | Cell Signaling Technology | 2690T |
| NF-κB p65 (D14E12) (1:1000) | Cell Signaling Technology | 8242S |
| Phospho-NF-κB p65 (Ser536) (93H1) (WB 1:1000; FC 1:1600) | Cell Signaling Technology | 3033S |
| Goat anti-Rabbit IgG (H+L) Cross-Adsorbed Secondary Antibody, Alexa Fluor 594 (1:2000) | Thermofisher Scientific | A-11012 |
| Phospho-IκBα (Ser32/36) (5A5) (1:1000) | Cell Signaling Technology | 9246S |
| IKBα (1:1000) | Cell Signaling Technology | 9242S |
| RIP3 (D4G2A) Rabbit mAb (1:500) | Cell Signaling Technology | 95702S |
| RIPk1 (D94C12) XP® Rabbit mAb (1:1000) | Cell Signaling Technology | 3493S |
| MLKL (D2I6N) Rabbit mAb (1:1000) | Cell Signaling Technology | 14993S |
| β-Actin (D6A8) Rabbit mAb (1:5000) | Cell Signaling Technology | 8457 |
| Anti-hTLR4-IgG (5 ug/ml) | InvivoGen | mabg-htlr4 |
| Mouse IgG1 Isotype Control | R&D Systems | MAB002 |
| Human TNF-alpha Antibody (10 µg/ml) | R&D Systems | AF-210-SP |
| Normal Goat IgG Control (10 µg/ml) | R&D Systems | AB-108-C |
| Human TNF RI/TNFRSF1A Antibody | R&D Systems | MAB225-SP |
| Human IL-10 R alpha Antibody (37607) (10 ug/ml) | R&D Systems | MAB274-100 |
| Human IFN-alpha/beta R2 Antibody (10 ug/ml) | R&D Systems | MAB4015-100 |
| **TLR ligands, inhibitors and recombinant proteins** | | |
| Recombinant Human IL-1 beta/IL-1F2 Protein | R&D Systems | 201-LB-005 |
| LPS | Invivogen | TLRL-3pelps |
| PAM3CYS | Invivogen | TLRL-PMS |
| Recombination Human CXCL4 | PEPROTECH | 300-16 |
| PF-4 (CXCL4) human | Sigma-Aldrich | SRP3142 |
| Brefeldin A Solution (1,000X) | Biolegend | 420601 |
| Z-VAD-FMK | R&D Systems | FMK001 |
| Necrosulfonamide | Sigma-Aldrich | 480073 |
| BMS-345541-IKKα/β inhibitor | Selleckchem | S8044 |
| Takinib | Selleckchem | S8663-5mg |
| RNase A | Sigma Aldrich | 10109142001 |
| DNase I | Sigma Aldrich | 10104159001 |
| SB202190, Hydrochloride, p38 inhibitor | Calbiochem | 559393 |
| CLI-095 TLR4 inhibitor | Invivogen | tlrl-cli95 |
| MRT67307 HCl (dual IKKϵ and TBK1 inhibitor) | Selleckchem | S7948 |
| TBK1/IKKε-IN-2 | MCE | HY-12453 |
| GSK8612-TBK1 inhibitor | Selleckchem | S8872 |
| SCH772984 | Selleckchem | S7101 |
| JNK Inhibitor II | Sigma Aldrich | 420119 |
| Magnolin | Selleckchem | S9102 |
